# Dual human milk oligosaccharide-fibre utilisation drives gut microbiome selection during weaning

**DOI:** 10.1101/2025.07.24.666491

**Authors:** Yunjeong So, Michael Jakob Pichler, Susanne Søndergaard Kappel, Chunsheng Jin, Carsten Eriksen, Ioanna Chatzigiannidou, Susann Teneberg, Karsten Kristiansen, Susanne Brix, Lise Aunsholt, Maher Abou Hachem

## Abstract

Gut microbiome (GM) maturation in early life follows organised taxonomic successions. How weaning impacts these trajectories remains underexplored. Here, we sampled faeces from seven mother-infant dyads at pre-, early and late weaning. Enrichment cultures (n=306) and metagenomic (n=108) analyses revealed an unexpected prevalence of fibre degradation genes and the growth of the pre-weaning infant GM on common dietary fibres. Utilisation of both human milk oligosaccharides (HMOs) and dietary fibres was revealed as a metabolic hallmark of the weaning GM. We showed that HMO-utilisation is retained beyond weaning, by analyses of maternal GM and HMO utilisation in 137 maternal isolates. Our findings highlight dual HMO-dietary fibre utilisation as a hitherto unrecognised driver that potentially orchestrates the selection of distinct adult GM species during weaning. This work outlines a plausible mechanism underlying the organised GM maturation in early life and highlights a previously overlooked role of HMOs during the weaning transition.

## Introduction

The gut microbiome (GM) assembly commences at birth^1,2^ and this community continues to mature into an adult-like structure during the first 1-3 years of life^3,4^. The mature GM structure is relatively resilient over time, hampering further colonisation of new members^5,6^. Perturbations in the assembly and maturation of the GM are associated with serious inflammatory^7,8^, metabolic^9,10,10,11^, and central nervous system disorders^12–14^. These aetiological links underscore the critical and persistent impacts of the early life GM on the developmental biology and health trajectory of the host^15^.

Diet is a key factor shaping the GM^1^. The onset of breastfeeding and weaning (solid food intake) represent critical nutritional milestones that elicit GM shifts. The most abundant component in mother’s milk, after lactose and fat, is a mixture of complex carbohydrates, collectively called human milk oligosaccharides (HMOs)^16^. HMOs are not digested by the infant, thereby serving as prime nutrients and selection cues for assembly of the neonatal GM^17^. Exclusively breastfed infants are predominantly colonised by HMO-utilising *Bifidobacterium* species^17^, whereas formula-feeding favours different communities^1,18,19^. During weaning, the infant’s diet gradually shifts from mother’s milk to solid food, rich in dietary fibres from grains, vegetables and fruits, marking a key phase in GM maturation. Besides HMOs, mucin or other host-derived glycans^20,21^ and dietary fibres are the main accessible nutrients for the weaning GM^22^. Maturation of the GM in early life follows defined taxonomic trajectories, despite high interindividual variations in microbial composition and dietary profiles^3,4^. The expansion of Bacillota is followed by Bacteroidota^1,2,23^, while the abundance of bifidobacterial HMO-utilising specialists declines. To date, breastfed infant cohorts are sampled by infant age^1,3,24–28^ and not nutritional status, precluding the assignment and matching of weaning nutritional profiles to the longitudinal metagenome sequence data.

Fluctuating HMO-dietary fibre gradients constitute a unique dietary signature during weaning, yet the mechanism by which this diet influences GM shifts remain underexplored. Here, we analysed the metagenomes of faecal samples and enriched communities from an infant-mother dyad cohort (Milkome cohort), designed specifically to monitor weaning progression. We investigated complex carbohydrate utilisation profiles of infant GM at three weaning stages, in addition to the maternal GM. Remarkably, we show that the ability to utilise dietary fibres is acquired before introduction of solid food, whereas HMO utilisation capability is maintained until late weaning and in the adult maternal GM. These findings unveil a mechanism for the selection of distinct core GM groups during weaning, which is orchestrated by the HMO-dietary fibre gradients, defining the weaning diet.

## Results

### Pre-weaning microbiome harbours both HMO and dietary fibre utilisation potential

To study the maturation of the infant GM during weaning, we established the Milkome infant-mother dyad cohort. Healthy mothers and their exclusively breastfed infants, who did not receive antibiotics or formula even during weaning, were recruited (Supplementary Table 1). Stool samples (n=18) from seven mothers and infants were collected at pre-, early and late weaning, defined as 100%, 70% (7 out of 10) and 30% (3 out of 10) of meals being mother’s milk, respectively, with the rest being solid food (Fig. 1a). For two dyads, either the late weaning or both the early and late weaning samples were not delivered (Fig. 1a). Freshly collected faecal samples were immediately equilibrated in an anaerobic chamber and enriched on selected carbohydrates as detailed below. We performed metagenomics shotgun sequencing of infant faecal samples from the three weaning stages, mother faecal samples collected at early weaning of their infants, and faecal consortia enriched on carbohydrates to allow strain-level annotation (Fig. 1b).

**Figure 1.**
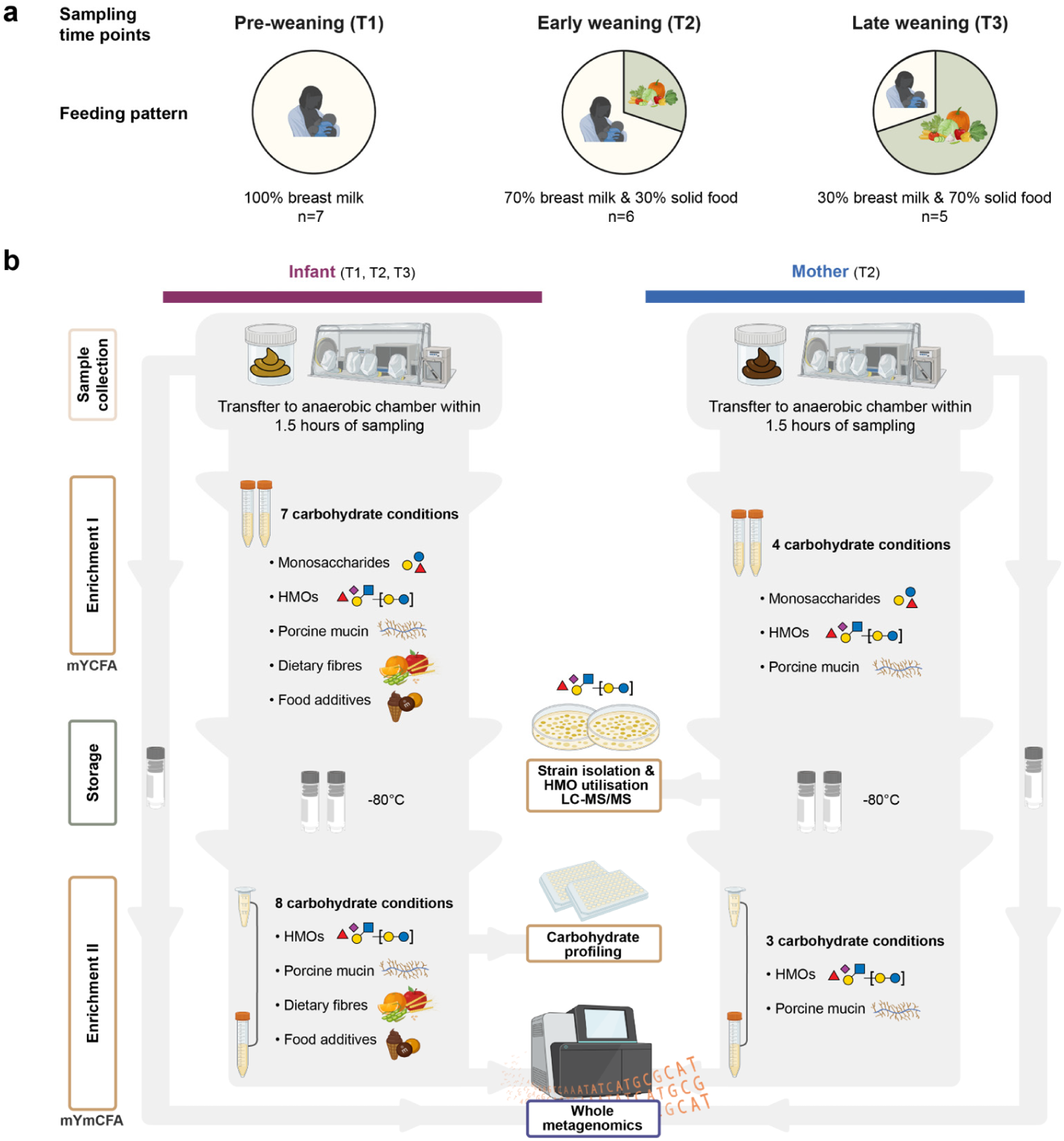
Sampling and analyses scheme of the Milkome cohort. **a** Cohort sampling coincided with the start of weaning and followed via changes in the distribution of mother’s milk to solid food meals. Pre-weaning (T1), early weaning (T2), and Late weaning (T3) sample collection was based on the specific proportion (%) of mother’s milk to solid food meals. **b** Sample collection, enrichment of faecal consortia on different carbohydrates, strain isolation, and whole metagenomic shotgun sequencing schemes.

As expected, Actinomycetota members displayed the highest relative abundances in 6 out of 7 infants in pre-weaning samples, with a trend of decreased abundance at late weaning (Fig. 2a). Low levels of Bacteriodota at pre-weaning, which increased in late weaning samples, concomitant with a reduction in Pseudomonadota, consistent with previously reported trends in early life^1,3,25^. The top abundant and prevalent *Bifidobacterium* species were key HMO-utilisers, *e.g.*, the specialist *Bifidobacterium longum* subsp. *infantis* (*B. infantis*)^17,29^, *Bifidobacterium longum* subsp. *longum* (*B. longum*)^30^ and *Bifidobacterium bifidum* that utilises mucin *O*-glycans and HMOs via extracellular glycosidases^31,32^. The abundance of *B. infantis* was at the cost of *B. longum* and the less prevalent *Bifidobacterium pseudocatenulatum,* both reported to utilise HMOs via different systems^33^ than *B. infantis*^33^ (Fig. 2b). These observations are consistent with the specialisation of *B. infantis* on HMOs, including the major fucosyllactose fraction (Supplementary Figs. 1a,b, and 2)^17,33^. *Bifidobacterium adolescentis*, a known plant fibre utiliser, not previously reported to utilise HMOs, was only observed in a single late weaning sample (I-B) (Fig. 2b), indicative of the poor competitiveness of non-HMO-utilising bifidobacteria, even at late weaning. Bacilli and Clostridia, besides Negativicutes, were also prevalent in the weaning samples but their HMO utilisation machinery remains underexplored. These data emphasise the important contribution of HMO utilisation for the competitiveness of bifidobacteria at least until late weaning.

**Figure 2.**
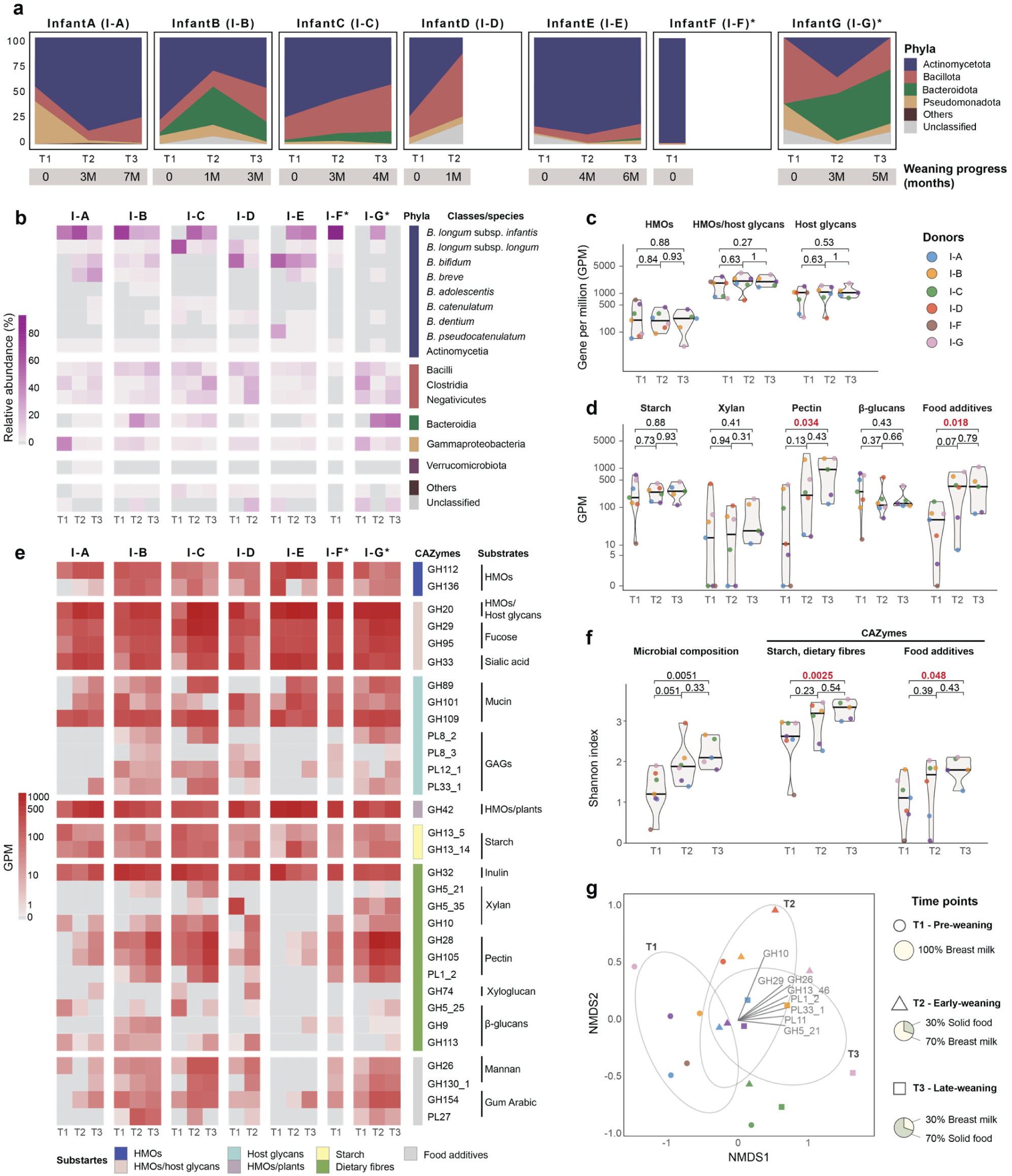
Complex carbohydrate-utilisation potential of infant microbiome during weaning. **a** Phylum level profiles of faecal bacteria at pre-weaning (T1, n=7), early-weaning (T2, n=6), and late-weaning (T3, n=5). Weaning progress in months illustrates the individual weaning rates. **b** Heatmap of species-level *Bifidobacterium* changes. Other taxa are depicted at class level. **c** Shifts in CAZymes targeting HMOs, HMOs/host glycans, and host glycans. **d** Shifts in CAZymes targeting starch, xylan backbone, pectin backbone, β-glucans including xyloglucans, and food additives. **e** Heatmap changes in different CAZyme families, organised by their carbohydrate substrates. **f** α**-**Diversity using Shannon index (vegan package^38^) across the weaning period, highlighting the diversification of species and CAZymes targeting starch, dietary fibres and food additives. P-values in panels **c** and **e** were calculated using the Wilcoxon test in R^39^. **g** Non-metric multidimensional scaling (NMDS) analysis of species composition, with an overlay of environmental fitting (envfit, in vegan package^38^) results, showing how selected CAZyme profiles correlate with species distribution. Selected CAZymes with significant correlations (P < 0.05) are shown. Distances between samples are calculated based on Bray-Curtis dissimilarity. *Infant whose mother had one dose of antibiotics during birth.

To analyse complex carbohydrate utilisation, we annotated carbohydrate-active enzyme (CAZyme)^34^ encoding genes in the metagenomic data. Specific CAZyme families and subfamilies were assigned as specific carbohydrate utilisation markers, based on biochemically characterised homologues (Supplementary Tables 2 and 3). Glycoside hydrolase (GH) families 112^35^ and 136^30,36^ that degrade the HMO type 1 building block lacto-*N*-biose (LNB) and the lacto-*N*-tetraose (LNT) backbone in HMOs, respectively, were selected as markers for HMO utilisation (Fig. 2c,d, Supplementary Figs. 1d and 2). Additionally, HMO-active β-galactosidases (GH42), fucosidases (GH29 and GH29), sialidases (GH33) and lacto-*N*-biosidases^34^ of GH20 were included (Supplementary Fig. 2, see Supplementary Results for details).

The potential to utilise HMOs was persistent during weaning, as indicated by the prevalence and abundance of genes encoding HMO-active CAZymes (Fig. 2c,e). By contrast, the utilisation potential of the host-derived glucosamine glycans (GAGs) appeared low, suggestive of the poor competitiveness of GAG degraders, *e.g.*, *Bacteroides* spp.^37^ at early weaning. Surprisingly, starch and dietary fibre utilisation genes were observed in all pre-weaning samples and their diversity increased during weaning (Fig. 2d-f). CAZymes targeting inulin, starch, xylan, pectin, and/or the food additive hydrocolloids β-mannan and gum Arabic were present already in pre-weaning samples (Fig. 2d-f), indicating that distinct fibre-utilising bacteria are acquired prior to introduction of solid food. Bacterial richness and evenness were only significantly different between pre-weaning and late weaning samples (Fig. 2e,f), consistent with the pivotal role of HMOs in shaping the GM, particularly at early weaning. This is also reflected by the abundance of HMO-utilising bifidobacteria, the abundance of HMO-utilisation genes, and the similar microbial community structure in the samples based on Bray-Curtis dissimilarity (R^2^ = 0.167, *P* = 0.0752) (Fig. 2b,e,g). Fibre (xylan, pectin, mannan)-specific CAZymes explained the segregation of distinct samples as weaning progressed (Fig. 2g), reflecting individual variations in GM composition and/or maturation rates during weaning.

### The weaning gut microbiome utilises both plant fibres and HMOs

Next, we sought to examine the carbohydrate utilisation profile by infant GM. We enriched freshly collected faecal samples on a mixture of monosaccharides (glucose/galactose/fucose), HMOs, mucin, and dietary fibres abundantly present in cereal grains (arabinoxylan, AX), vegetables and fruits (xyloglucan, XC; pectin, PEC) (Supplementary Figs. 3,4 and Supplementary Tables 4-6) as well as in natural food additives (galactomannan, GMN; gum Arabic, GA). These enrichments were performed twice (enrichments I and II) (Fig. 1) to allow rapid handling of the fresh faecal samples by using fibre blends in the first enrichment. The second enrichment was performed using an optimised medium (Supplementary Table 4) on individual complex carbohydrate substrates.

Infant faecal consortia from both enrichments grew on HMOs (Fig. 3a,b and Supplementary Fig. 5). Growth on mucin in pre-weaning samples was weak or insignificant (Fig. 3b). Interestingly, broad growth on dietary fibres was observed in pre-weaning samples, especially for AX, followed by PEC/GMN and XG/GA (Fig. 3b, Supplementary Figs. 5 and 6a,b). This was in line with growth on AX and PEC by all infant early and late weaning samples (Fig. 3b and Supplementary Fig. 5). Growth of pre-weaning infant GM on dietary fibres has to our knowledge, hitherto, not been demonstrated.

**Figure 3.**
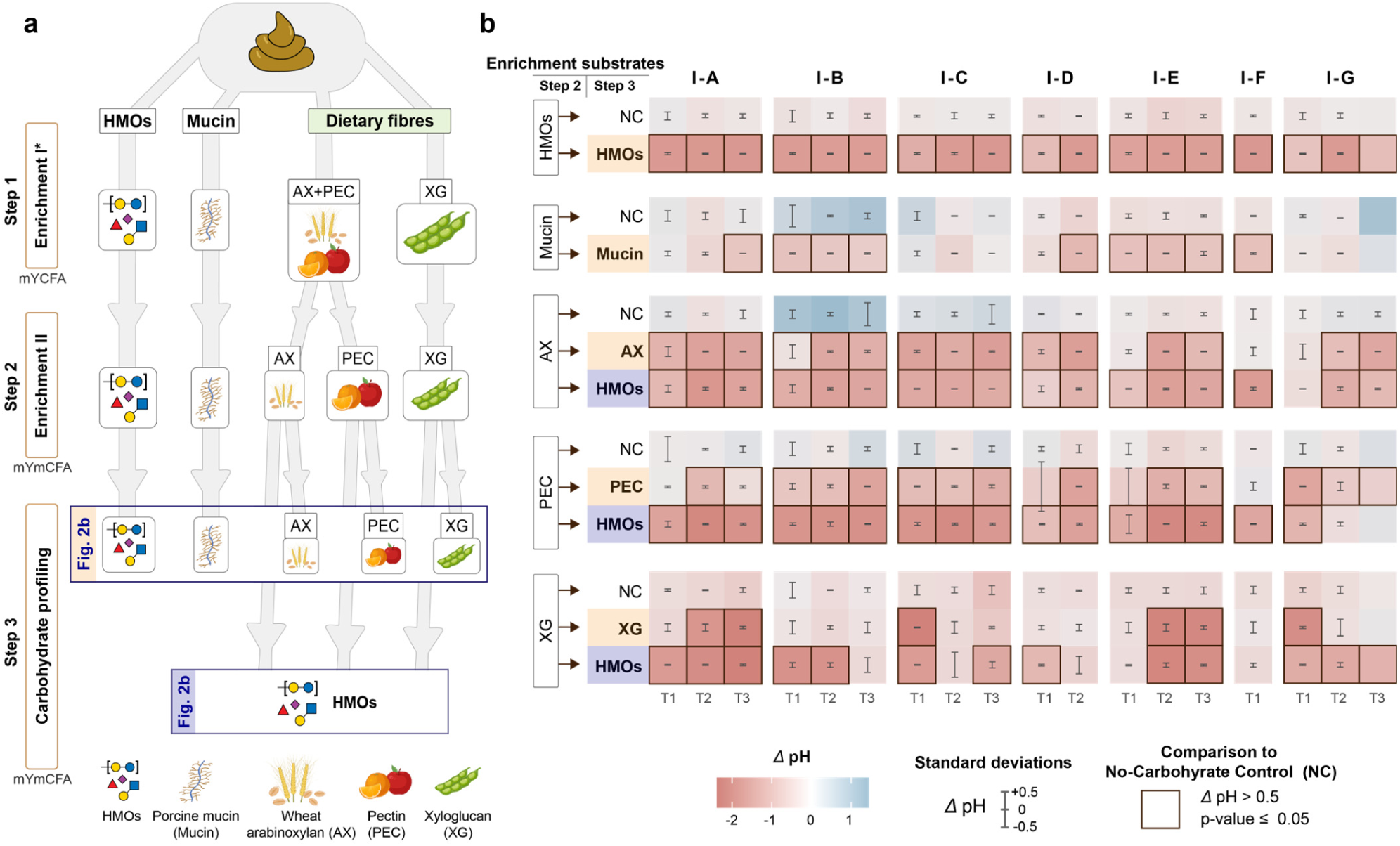
Growth of faecal consortia on HMOs, mucin and dietary fibres. **a** Workflow for faecal sample enrichment. Enrichment I was performed on yeast extract-reduced YCFA medium (mYCFA), whereas enrichment II was conducted in a further optimised medium (mYmCFA), with reduced yeast extract and protein (Casitone) content. **b** Changes of pH after a 24-hour enrichment on HMOs, porcine mucin, and dietary fibres. Dietary fibres consortia were subsequently re-grown on HMOs to evaluate dual HMO-fibre utilisation capacity. Error bars depict standard deviations of three technical replicates. The *P*-values were calculated using a t-test with two-tailed distribution with unequal variance. Individual *P*-values provided in the source data file.

We then selected a Swedish mother-infant dyad cohort (n=98)^11^ with available nutritional^40^ and metagenomic data to investigate the prevalence of potential growth on fibres before weaning. We annotated the fibre degradation potential in exclusively breastfed infants sampled at the first week (n=53) and 4^th^ month of life (n=51), similarly to above. All analysed metagenomes (n=104) harbour the potential to utilise ≥1 fibre type (Supplementary Fig. 7), in agreement with findings from our cohort. Despite interindividual variations, the median abundances of the selected CAZyme genes indicated a considerable fibre utilisation potential of the GM during exclusively breastfeeding (Supplementary Fig. 8). The demonstrated fibre utilisation by preweaning faecal consortia and metagenomic analyses of both cohorts are consistent with the presence of distinct fibre utilising species in the GM of breastfed infants prior to weaning. The poor growth on mucin suggested that HMOs may sustain fibre-utiliser during exclusive breastfeeding. To test this, we examined the growth of the fibre-enriched consortia (enrichment II) on HMOs. Interestingly, most consortia displayed growth on HMOs at all sampled weaning stages (Fig. 3a,b, Supplementary Figs. 5 and 6b). These findings unravel the dual ability to utilise both HMOs and dietary fibres as a hitherto undescribed signature of the weaning GM.

### Genes encoding HMO degradation during weaning

We explored the composition of the enriched consortia from the infant faecal samples on both HMOs and mucin. The dominance of *B. infantis* in HMO enrichment cultures, except for one infant (I-E), was similar or higher than the corresponding faecal samples (Fig. 4a). This and the considerably lower relative abundance on mucin, reflect the specialisation of *B. infantis* on HMOs (Fig. 4a). By comparison, the preference of *B. bifidum* towards mucin *O*-glycans was evident in enrichments on mucin. Interestingly, *Enterococcus* species were relatively abundant in HMO enrichments, especially in pre-weaning samples, together with other Bacilli and Clostridia (*viz*. *Clostridium* and *Ruminococcus* spp.), indicating that distinct members of these taxa are endowed with the machinery to assimilate HMOs or their transformation products. Overlap in growth on HMOs and mucin was observed for *Enterococcus* species, potentially suggesting a less discriminative enzymatic machinery. Key butyrate producers^41^, *e.g.*, *Roseburia* (I-A,) and *Faecalibacterium* (I-B), increased in abundance in the late weaning mucin enrichment, consistent with their ability to cross-feed on mucolytic bacteria^20^.

**Figure 4.**
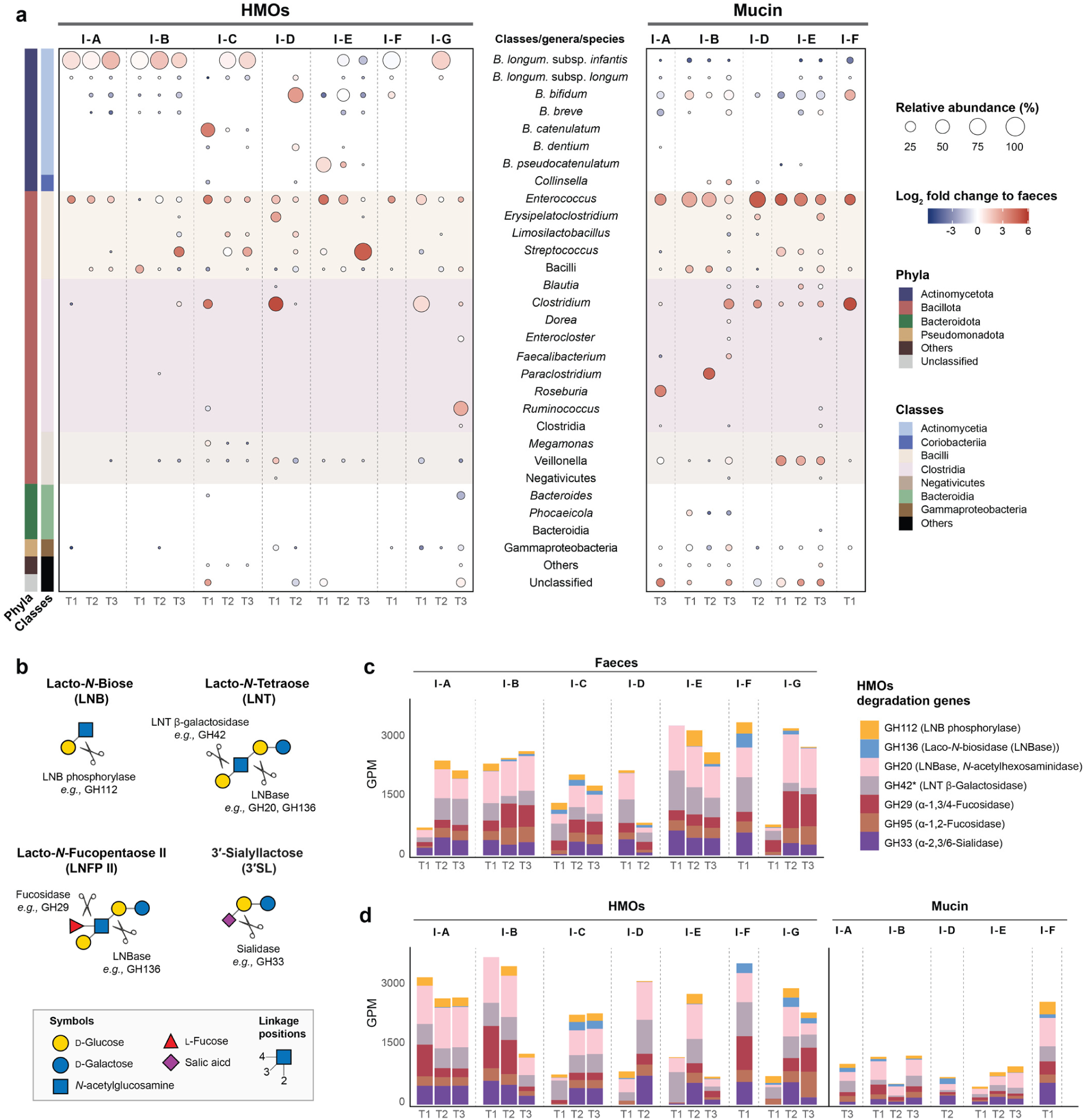
Infant faeces consortia enriched on HMOs and Mucin. **a** Enriched microbial community composition. **b** Schemetic representation of selected enzymatic activities targeting common HMO motifs. **c** and **d** Abundance of genes conferring the degradation of HMOs in original faeces and consortia enriched on HMOs and mucin. Gene counts are normalised as gene per million (GPM).

We also compared genes encoding putative HMO backbone-degrading enzymes (GH112, GH136), sialidases and fucosidases (GH33, GH29, GH95), as well as hexosaminidases and less-common lacto-*N*-biosidases, both from GH20 (Fig. 4b-d, Supplementary Fig. 2). The selected genes were abundant in most samples, underscoring a pivotal role of HMOs in GM selection even at late weaning (Fig. 4c,d). The markedly lower abundance of HMO degradation genes in mucin enriched consortia validated the preferential roles of the selected CAZymes in HMO utilisation during weaning (Fig. 4d).

### Fibre utilisation gene signatures in the weaning infant GM

We analysed the composition of the fibre-enriched consortia to explore the taxonomic groups and fibre degradation genes during weaning. As expected, *B. infantis*, was only observed in a single early weaning enrichment on AX (Fig. 5a), consistent with the inability of most strains to utilise fibres. By contrast, *B. longum* increased in abundance on AX and on GA relative to the faecal samples (see early weaning, samples A, B and E) (Fig. 5a and Supplementary Fig. 6c). This trend also applied to *B. breve* after enrichment on AX and PEC in pre-weaning samples (Fig. 5a). This evidence supports the growth of distinct HMO-utilising bifidobacteria on plant fibres. Moreover, Bacilli, especially *Enterococcus* species were also enriched on plant fibres (Fig. 5a and Supplementary Fig. 6c). Interestingly, the relative abundance of several Clostridia increased in the fibre enrichments, relative to the faecal samples, including two *Clostridium* species and the butyrogenic *Roseburia intestinalis* (Fig. 5a), a primary degrader of xylan and mannan^42,43^.

**Figure 5.**
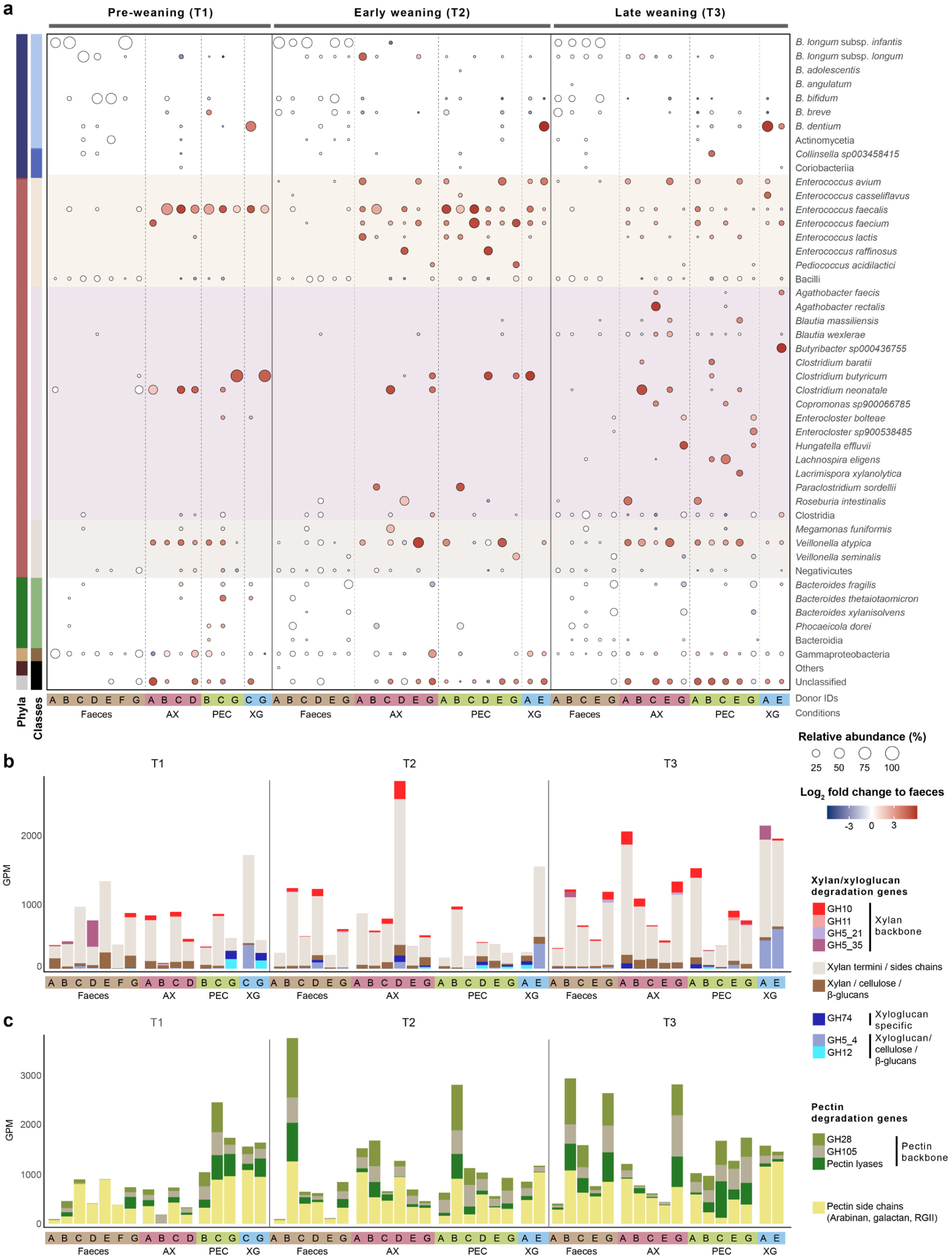
Infant faeces and faeces consortia enriched on dietary fibres, arabinoxylan, pectin, and xyloglucan. **a** Microbial community composition in faecal samples and enriched consortia based on metagenomic analyses. **b** Abundance of selected genes mediating the degradation of xylan (AX) and xyloglucan (XG) in faecal samples as well as arabinoxylan and xyloglucan-enriched consortia. **c** Abundance of genes mediating pectin (PEC) degradation in the faecal samples and enriched consortia. In panels **b** and **c**, the normalised gene abundance is depicted as gene per million (GPM).

Next, we sought to rationalise the enrichment profiles on the selected fibres. Genes encoding fibre-specific CAZymes were identified in both the faecal samples and fibre-enriched consortia (Fig. 5b,c and Supplementary Fig. 6d-f). The enrichment increased the abundance of endo-acting CAZymes critical for polysaccharide backbone degradation, *e.g.*, xylanases, pectin lyases and mannanases on target fibres (Fig. 5b,c and Supplementary Fig. 4d,f). Interestingly, some of these CAZymes were only observed after enrichment, but not in the original faecal samples (Fig. 5b,c, Supplementary Results), underscoring their pivotal roles for polysaccharide utilisation. Increases of CAZyme genes on non-substrate fibres, *e.g.*, increase in xylanase genes in consortia enriched on pectin, indicate that distinct primary degraders possess CAZyme arsenals to degrade multiple fibres. Our metagenomics and culture enrichment analyses demonstrate that fibre-utilising GM species are already present during exclusive breastfeeding, hinging primarily on their capacity to utilise HMOs. Expansion of these species and further selection of bacteria with similar catabolic profiles, is promoted as weaning progresses.

### Maternal GM grows on HMOs

The proliferation of dual HMO-fibre utilisers during weaning implies that HMO utilisation pathways may be retained beyond GM maturation. To investigate this, we enriched the maternal faecal samples on HMOs. Enrichments on mucin aimed at assessing if growth on HMOs is driven by mucin degraders, due to structural overlap of glycans from both substrates. All faecal samples sustained good growth on HMOs. Surprisingly, significant growth on mucin was only observed in two samples (Fig. 6a). As expected, the maternal faecal GM (R^2^ = 0.028, *P* = 0.0003) and the enriched consortia on HMO (R^2^ = 0.018, *P* = 0.0056) segregated from infant samples (Supplementary Fig. 9a). High abundance of Bacillota and Bacteriodota and low Actinomycetota abundance was observed as opposed to the infant samples (Fig. 2a, Supplementary Fig. 9b). The *Bacteroides* genus was abundant in all mothers but was depleted after enrichments on HMOs and was markedly reduced in the mucin cultures (Fig. 6b, Supplementary Fig. 9b). A considerable increase of Actinomycetota, was observed in the HMO cultures, mainly attributed to *Bifidobacterium* HMO utilisers, *e.g.*, *B. pseudocatenulatum*, *B. catenulatum* and *B. longum*. Bacilli and Clostridia, were also enriched on HMOs. Abundance of *B. bifidum* increased in consortia enriched on HMOs and mucin in two samples (mothers D and E) (Fig. 6b). Generally, the enrichment of species was specific to either HMOs or mucin, consistent with our infant analyses. These findings underscore that specific adult GM are adapted to growth on HMOs but not conjugated mucin *O*-glycans. Bacteroidota were not competitive on HMOs, consistent with their delayed expansion until solid food becomes the prime GM nutrient.

**Figure 6.**
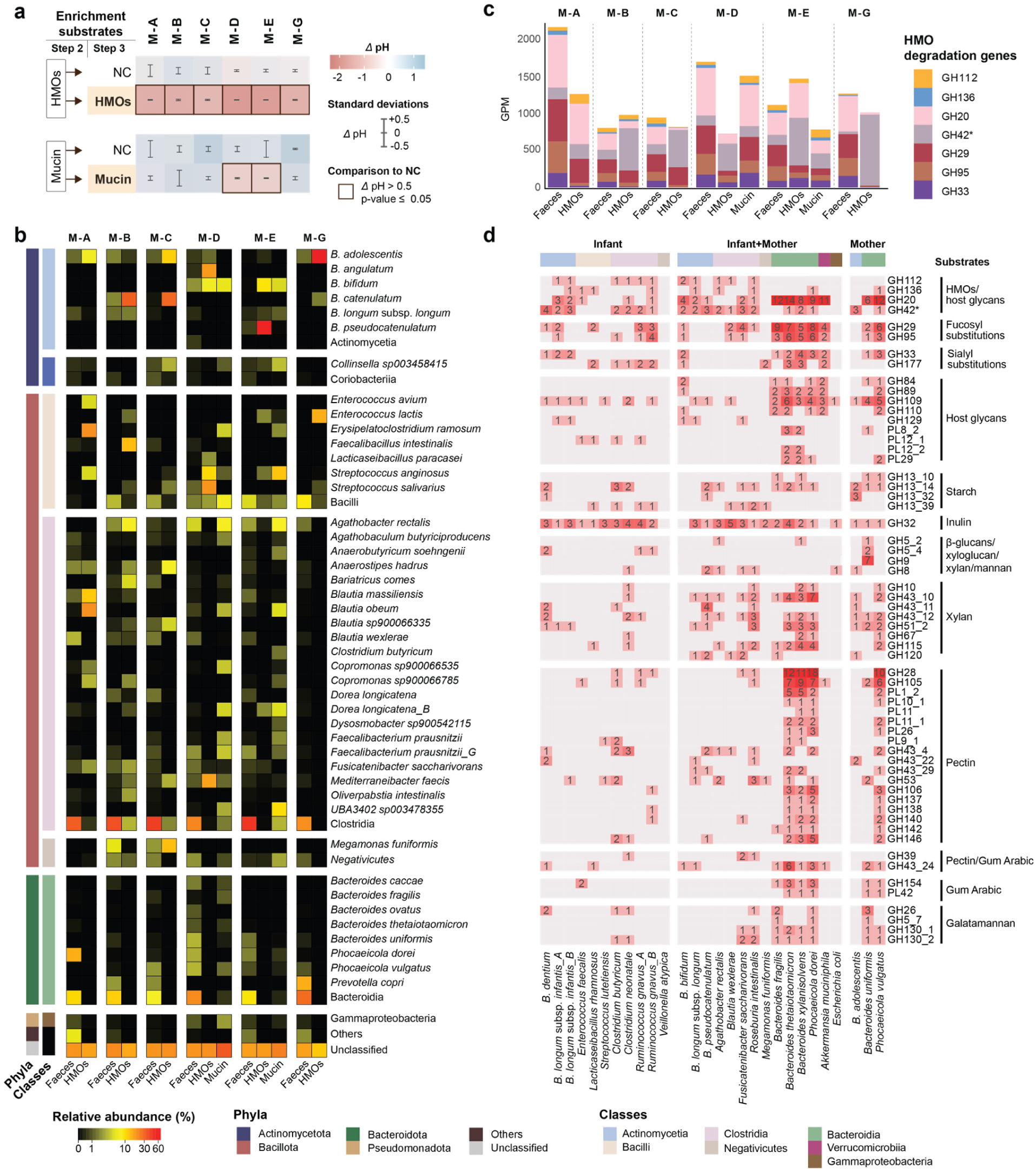
Maternal faeces and faecal consortia enriched on HMOs and mucin. **a** Changes of pH after a 24-hour enrichment on HMOs and mucin, *P*-values are calculated with a t-test. **b** Microbial community composition in faecal samples and enriched consortia based on metagenomic analyses. **c** Abundance of HMOs degradation genes in faecal samples and consortia enriched on HMOs and mucin. The normalised gene abundance is depicted as gene per million (GPM)**. d** Metagenome assembled genomes (MAGs), retrieved from both infant and maternal faeces as well as the enriched consortia, were selected based on the top 25 most abundant in each infant and maternal faeces. *Clostridium butyricum* was additionally included due to its high abundance in the enriched consortia. CAZyme families with more than three gene copies across the selected MAGs are shown and classified based on target substrate.

Then, we analysed genes encoding HMO-degradation by maternal GM. Growth on HMOs enriched β-galactosidase (GH42) genes, whereas counterparts encoding GH112 displaying dual activity on the HMO I block (LNB) and the mucin-derived galacto-*N*-biose (GNB, Gal-β1,3-GalNAc) featured in consortia enriched on both HMO and mucin. Overall, different CAZyme profiles were associated with growth on HMOs and mucin, reiterating the presence of HMO-specific CAZymes (Fig. 6c). For a strain level view, we analysed the 256 high quality MAGs, retrieved from infant and mother faeces as well as enriched consortia. Two MAGs from *B. infantis* (infant samples) and one *B. bifidum* encode mainly HMO-active enzymes (Fig. 6d and Supplementary Fig. 10) but possess 1-4 plant fibres encoding genes (GH42, GH32, GH51_2, GH53 and/or GH43_24). Only few infant consortia enriched on dietary fibres contained *B. infantis* and *B. bifidum* at reduced abundance compared to the faecal samples (Fig. 4a), suggesting limited competitiveness of both species on plant fibres. By contrast, *B. longum* and *B. pseudocatenulatum* were found both in infant and mother samples. Both species encode HMO and plant fibre utilisation enzymes (Fig. 6d, Supplementary Fig. 10), consistent with their enrichment on both these substrate types. The remaining *Bifidobacterium* MAGs showed mainly plant fibre utilisation potential, besides the presence of one or multiple GH42 genes, which may target either plant fibres or HMOs, *e.g.*, LNT, (Fig. 6d, Supplementary Figs. 9c and 10b). The MAGs of butyrate producers in maternal and infant samples, *e.g., Agathobacter rectalis* (*Eubacterium rectalis*) and *R. intestinalis,* encode both plant-derived fibres and HMOs/mucin utilisation genes. (Fig. 6d and Supplementary Fig. 10b,c). In summary, our work unravels the growth of maternal GM bacteria on both HMO and dietary fibres, consistent with the metagenomic CAZyme analyses.

To further investigate HMO utilisation by adult GM, we isolated 258 strains from maternal faecal consortia, enriched on HMOs. Regrowth and replating resulted in 188 isolates showing significant growth (*P* < 0.01) relative to no-carbohydrate controls.

To elucidate HMO utilisation preferences, we grew 137 isolates on 1% (w/v) HMOs at 37 °C for 24 h (Supplementary Figs. 11 and 12a,b). The isolates clustered into five HMO groups, based on their HMO depletion profiles on 22 selected HMOs (Fig. 7, Supplementary Figs. 12c, and 13, and Supplementary Results). Group I had the highest growth median, which was associated with the depletion of abundant undecorated LNT/LN*n*T (lacto-*N*-*neo*tetraose) HMO backbones (Supplementary Fig. 1), including those branched with β1,6-*N*-acetylglucosamine unit (Supplementary Fig. 13). By contrast, groups II and III lacked activity on LNT/LN*n*T, but removed α2,3- and/or α2,6-linked sialic acid (Neu5Ac) caps, correlating to lower growth median than group I (Supplementary Fig. 12c). In summary, the depletion of common HMO mainchains and their plausible assimilation, correlates to efficient growth by the maternal isolates.

**Supplementary Data Fig. 8.**
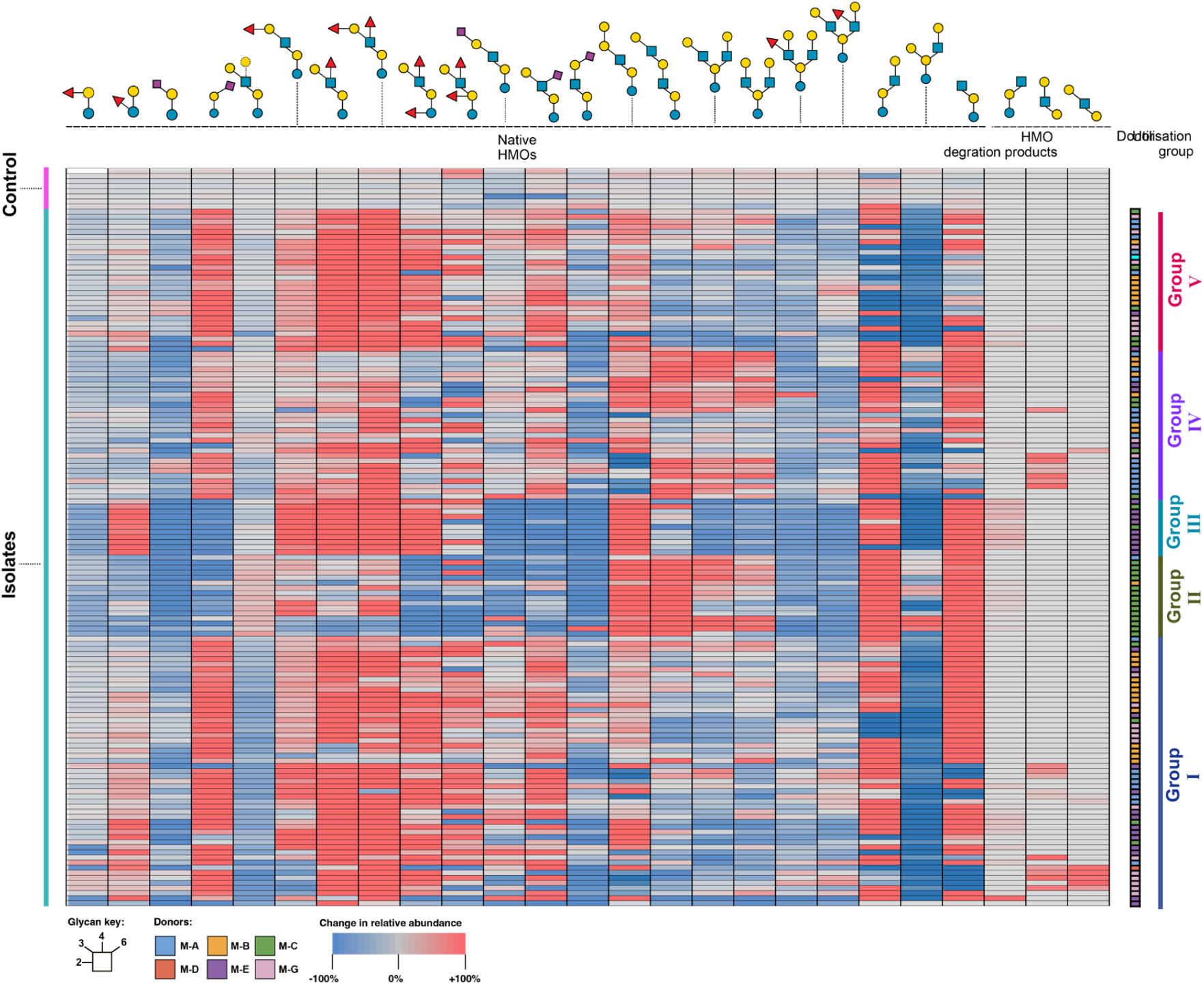
HMO utilisation profiles of adult gut bacteria. Heatmap showing percentage changes in the relative abundance of 22 selected HMO structures and three potential HMO degradation products by individual isolates compared to uninoculated medium controls. Changes in relative abundance profiles of HMO structure were analysed and clustered as a metric for similarities in utilisation and/or enzymatic modification among the isolates. Hierarchical clustering was performed using the “group average” clustering method with Euclidean distance as the distance measure using OriginPro 2023.

The HMO-degrading genes (GH112 and GH136) were identified in bifidobacteria, but also in several Clostridia from the maternal GM. To assess the broad relevance of these observations, we analysed the taxonomic distribution of both CAZyme families in the entire human GM catalogue^44^. Surprisingly, Actinomycetota (including *Bifidobacterium*) harboured the minority of GH112/GH136 sequences (Supplementary Fig. 14). Most GH112 (84%) and GH136 (60%) sequences were from Bacillota, consistent with the growth of *Roseburia* and *Eubacterium* on HMOs, mediated by GH112 and GH136 enzymes^36^. The presence of these two enzyme families in Eubacteriales (*e.g.*, *Ruminococcus*) and Bacilli (*e.g.*, enterococci) may rationalise the enrichment of both taxonomic groups on HMOs (Fig. 4a, Supplementary Fig. 14, and Supplementary Table 7). Multi-sequence alignments and phylogenetic analyses further supported the preponderance of GH112/GH1136 CAZymes in core GM species, especially Bacillota (Supplementary Fig. 15a-d). Finally, the conservation of active site residues in both families is suggestive of a high selection pressure to maintain their function (Supplementary Fig. 15e,f). Altogether, these findings highlight a previously not recognised prevalence of HMO utilisation enzymes in bacteria, considered as core adult GM species.

## Discussion

Breastfeeding for at least 1-1.5 years has been common until shorter lactation was inferred in the 19^th^ century Western societies, ramping up child mortality rates^45^. The World Health Organization (WHO, https://www.who.int) recommends at least 6 months of exclusive breastfeeding, followed by a weaning period up to 2 years of age^46^, but less than half of the infants under 6 months are breastfed according to WHO.

This study focuses uniquely on weaning by sampling three different stages, with pre-weaning as a reference. We demonstrate that fibre-utilising bacteria colonise infant guts before introduction of solid food. This notion is supported by the conservation of fibre utilisation genes in pre-weaning infant GM in our cohort and a Swedish infant cohort^1^, as well as growth of pre-weaning faecal consortia on dietary fibres. The ability of specific fibre-utilisers to additionally harness HMOs as nutrients, albeit less efficiently than the specialists like *B. infantis*, allows for their persistence at low abundance during exclusive breastfeeding. Weaning triggers a gradual and organised expansion of dual HMO-fibre utilisers, commensurate with the increase in solid food to mother’s milk ratio. This strategy has likely evolved to select for core GM species and suppresses random colonisation by environmental microbes during the critical maturation transition. At weaning completion, the gut mucosal surfaces are densely colonised by a core GM community, which allows for more liberal and diet-driven diversification of infant GM. This proposed mechanism is in line with the dominance of HMO-utilising bifidobacteria in late weaning samples, which is key to the maturation of the infant immune system, *e.g.*, via aromatic lactate metabolite secretion, *e.g.*, by *B. infantis*^47^. Despite the abundance of HMOs in weaning diet, hitherto the maturation of the GM has been largely attributed to the introduction of solid food^48–50^.

At present, Bifidobacteria are classified either as adult or infant species and focus has been either on their HMO or fibre utilisation. Recently, utilisation of both HMOs and plant-derived fibres was reported in *Bifidobacterium catenulatum* subspecies *kashiwanohense*^51^, having a low prevalence in both infants and adults^51^. Similarly, a *B. longum* clade similarly endowed to utilise both HMOs and plant fibres was shown to be prevalent in weaning infants Bangladesh, but not other geographic locations^52^. Here we show that dual HMO-plant fibre utilisation applies to other bifidobacteria, *e.g.*, *B. breve* and *B. longum*, which possess enzymes targeting both components of the weaning diet^30,53^. A hitherto not reported finding is the broad taxonomic distribution of HMO type 1 utilisation markers in core GM groups other than *Bifidobacterium* species. Previously, we reported that primary fibre degraders from *Roseburia* and *Eubacterium* grow on abundant HMO type 1 backbones^36^. Cross-feeding on sialic acid form HMO or mucin degradation maybe be mediated by a sialic acid (Neu5Ac) importer conserved in Clostridial GM members, including *Ruminococcus*^54^ and *Faecalibacterium* species, which showed increased relative abundance in the mucin enrichment. Increased butyrate production by early colonising Clostridia during weaning^55^ has been proposed to be important for immune modulation and protection from immune pathologies. By comparison, the poor competitiveness of *Bacteriodes* glycan utilisation generalists during growth on HMOs, is consistent with their delayed expansion towards cessation of breastfeeding, as diverse solid foods dominate the infant nutrition.

In conclusion, this study highlights the role of HMOs in the maturation of infant GM during weaning. We reveal the presence of dual HMO and fibre-utilising bacteria before the introduction of a solid food diet. This core weaning GM gradually blossoms and expands following the onset of weaning. The ability to utilise HMOs (specifically type 1 blocks) besides diverse fibres common in solid food, is likely to be a critical selection criterion that underpins the organised colonisation and proliferation of core GM taxa during the weaning transition. This orchestrated maturation is likely to have important implications on the developmental biology and health outcomes of the host. A consequence of this previously unrecognised evolutionary adaptation is the persistence of the HMO utilisation trait in adult GM, as demonstrated in this study. Further research is required to shed light on the weaning transition to study how timing and rate of weaning affect GM maturation and potentially the health profile of the host.

## Methods and materials

### The Milkome Cohort

The study, centred on the Milkome cohort, was approved by the Ethical Committee of the Capital Region of Copenhagen, Denmark (study registration ID H-21067700) and registered at Clinicaltrials (https://clinicaltrials.gov, registration ID NCT07026526). The cohort comprises seven mother-infant dyads who were recruited at Copenhagen University Hospital, Rigshospitalet (Copenhagen, Denmark) between August 2022 and January 2024, based on informed consent obtained from the mothers (maternal consent) and the parents (infant consent). Detailed information about volunteer inclusion and exclusion criteria are summarised in Supplementary Table 1.

### Sample collection

Infant and maternal faeces were collected by participants at three distinct weaning stages: Pre-weaning (exclusive breastfeeding), early weaning (seven out of ten consecutive meals given to the infant within a period of 1-2 days, would be mother’s milk and three would be solid food), late weaning (three out of ten consecutive meals given to the infant within a period of 1-2 days would be mother’s milk and seven would be solid food). An overview of the sample collection scheme and analyses is reported in Fig. 1. Faecal sample collection was designed to maximise viability and recovery of the GM via 1) preserving an anaerobic headspace in sample containers, 2) cold storage and minimal time (<1.5 h) from collection to laboratory delivery, and 3) enrichment of faecal consortia from the freshly received samples. Faecal samples were collected using a commercial anaerobic microbiome collection kit (GutAlive, Gijón Spain) and processed within 1.5 h of collection in an anaerobic chamber (Don Whitley, Bingley, UK or Coy Laboratory Products, Grass Lake, MI, US). Upon arrival, samples were equilibrated anaerobically at 37°C for 30 min and thoroughly homogenised. One 0.5 g aliquot was immediately processed (*vide infra*), while the remaining aliquots were stored in cryotubes at −80°C.

### Preparation of Human milk oligosaccharides (HMOs) and porcine gastric mucin growth substrates

Human milk oligosaccharides (HMOs) were purified from pooled human milk samples, (purchased or donated from Hvidovre hospital, Hvidovre, Denmark), by solid phase extraction (SPE) as previously described^56^. In short, milk fat was removed by centrifugation (10.000 *g*, 30 min, 4°C), and proteins were precipitated by the addition of 3 volumes of ice-cold ethanol (99%) and incubation for 18 h at 4°C. Lactose in the protein free supernatants was digested with a commercial ß-galactosidase from *Kluyvermomyces lactis* (Sigma Aldrich) (20 U mL^−1^, 3 h at 37 °C). Then, residual lactose and monosaccharides were removed by SPE using an in-house column packed with a 100 g of activated charcoal (20-40 mesh particle size, Sigma Aldrich). The column was activated with one column volume (CV) of 80 % (v/v) acetonitrile (ACN) in milliQ water and equilibrated with two CVs of 4 % (v/v) ACN in milliQ water (solution A), before samples were loaded. Residual lactose and monosaccharides were removed by washing with six CVs of solution A and HMOs were eluted with two CVs of 40 % (v/v) ACN in water. A rotary evaporator was used to remove residual ACN, before the HMOs were freeze-dried. The residual lactose concentration in the eluted HMOs was determined by high-performance anion exchange chromatography with pulsed amperometric detection (HPAEC-PAD). For this, samples (10 µL) were injected into a 3 × 250 mm CarboPac PA200 column (Thermo Fisher Scientific) and a 3 × 50 mm CarboPac PA200 guard column (Thermo Fisher Scientific) installed on an ICS-5000 (Dionex) system for the separation. The bound saccharides were eluted with a stepwise linear gradient of sodium acetate: 0-7.5 min of 0-50 mM, 7.5-25 min of 50-150 mM and 25-35 min of 150-400 mM, at a flow rate of 0.35 mL min^−1^ and a mobile phase of constant 0.1 mM NaOH. Standards (0.01-0.5 mM) of lactose in milliQ water were used to quantify residual lactose as described above. The analysis was performed in independent triplicates. The lactose-depleted HMO preparation (0.42 ±0.02 % (w/w) lactose in the HMO preparation, corresponding to 61 µM lactose in standard growth medium containing 0.5% (w/v) HMOs, was stored at −20°C until use for bacterial growth, enrichment and isolation experiments., which was deemed insufficient to markedly impact the growth outcomes.

Porcine gastric mucin type III (PGM) was further purified as previously described^57^. In short, mucin was dissolved to a final concentration of 2.5 % (w/v) in phosphate buffered saline, pH 7.4 (PBS) and stirred for 20 h at room temperature. Insoluble mucins were removed by centrifugation (10.000 *g*, 30 min, 4 °C) and soluble mucins were precipitated from the supernatants by adding 3 volumes of ice-cold ethanol (99 % v/v) for 18 h at 4°C. Next, purified soluble mucin was dialysed in milliQ using a SpectraPor^TM^ 50 kDa molecular weight cut-off membrane (Fisher Scientific, Waltham, MA, USA), freeze-dried, and stored at −20 °C.

### Enrichment of faecal consortia on selected carbohydrate sources

Initially we tested the YCFA medium on the first received samples, however, high growth on the no carbohydrate-supplemented control prompted us modify the medium to reduce baseline growth. To this end, yeast extract content was reduced by 50% in the first enrichment (mYCFA). Fresh faecal samples (0.5 g) from mothers and infants were diluted (1:10 w/v) in mYCFA medium^58^ (0.1% (w/v) yeast, 1 % (w/v) Casitone and 0.05 % (w/v) PGM (Supplementary Table 3) and thoroughly homogenised for 5 min using a vortex mixer. This medium was additionally designed to mimic the *in vivo* presence of mucin, which may serve as an adhesion surface for distinct GM groups. Following sedimentation of insoluble particles, 400 µL of supernatant was transferred to a fresh tube and subsequently diluted three additional times 1:10 (v/v), resulting in a final dilution of 10^-3^ in mYCFA. The diluted samples were used to inoculate 6.5 mL mYCFA medium, supplemented with 0.5 % (w/v) carbohydrates (Supplementary Table 4) sterilised by filtration (0.5 µm filters) or standard autoclaving (for insoluble carbohydrates), to a final sample dilution of 10^-5^.

The cultures were incubated for 24 hours at 37 °C in an anaerobic chamber (Don Whitley, Bingley, UK or Coy Laboratory Products, Grass Lake, MI, US) and bacterial growth was monitored by measuring *OD*_600_ and pH. Cultures were performed in technical duplicates. Enriched bacterial consortia were supplemented with glycerol to a final concentration of 25% (v/v) and immediately frozen at −80°C.

To further optimise the specificity of the carbohydrate enrichments and reduce potential protein-mediated growth by proteolytic bacteria, the enriched consortia (*vide supra*) were subjected to an additional enrichment round on the same carbohydrate sources, using a reduced Casitone content (0.5% w/v) medium (mYmCFA) (enrichment II, Supplementary Tables 6 and 4). For this, pre-cultures were prepared by inoculating 1 mL mYmCFA medium containing 0.5% (w/v) corresponding carbohydrate (1% (w/v) for HMOs), with 40 µL of each cryopreserved enriched consortium (Fig. 1b). Pre-cultures were grown to mid-late exponential phase (*OD*_600_ 0.5-0.9) and then used to inoculate cultures for two experimental setups: a) metagenomics shotgun sequencing and b) carbohydrate utilisation profiles. For metagenomic shotgun sequencing, cultures were inoculated in either 10 mL of fresh mYmCFA with 1% (w/v) HMOs or 20 mL of fresh mYmCFA with 0.5% (w/v) mucin or polysaccharides to accommodate the lower growth mucin or polysaccharides (single experiment). These cultures were harvested after 24 h, except for those supplemented with HMOs, which were harvested after 8 h, due to more rapid growth. Following centrifugation (4,000 *g*, 15 min, 4°C) cells were washed twice with ice-cold sterile PBS and then stored at −80°C for DNA extraction. For carbohydrate utilisation profiling (Fig. 2), cultures were inoculated into 96-well microtiter plates in 280 µL of fresh mYmCFA containing 0.5% (w/v) carbohydrates (1% (w/v) for HMOs) in triplicates, each to a starting *OD*_600_=0.05. Growth was monitored by *OD*_600_ using a BioTek Epoch 2 microtiter plate reader (Agilent, Santa Clara, CA, USA) and pH measurements after 24 hours, which were then compared to non-inoculated media.

### Isolation and growth of bacterial strains

For isolation, HMO enriched consortia from maternal samples (sample: early weaning of the infant) were grown in 2 mL mYmCFA supplemented with 1% (w/v) HMOs to late exponential phase (*OD*_600_ 0.7-0.9). Next, the enriched cultures were serially diluted 200-fold in max (1:10 steps (v/v)) in sterile mYmCFA to a final dilution of 10^-3^ and thereafter spread (20 µL) on mYmCFA agar plates containing 1% (w/v) HMOs. Plates were incubated anaerobically at 37°C for 2-3 days. From each maternal sample, 50 single colonies were randomly picked and re-streaked twice on a fresh mYmCFA, supplemented with 1% (w/v) HMOs agar plate. The isolates were then transferred to liquid mYmCFA media supplemented with 0.5 % (w/v) glucose, 0.5 % (w/v) lactose and 0.5 % (w/v) maltose, grown for 18 h, and stored at −80°C in a 25 % (v/v) glycerol suspension. The enrichments were performed in a single experiment. To explore the HMO utilisation capacity, cryopreserved isolates were grown in 1 mL mYmCFA supplemented with 0.5 % (w/v) each of glucose, lactose and maltose to mid-late exponential phase (*OD*_600_ 0.5-0.7). These pre-cultures were used to inoculate 200 µL mYmCFA medium with 1 % (w/v) HMOs to a starting of *OD*_600_ of 0.05 and growth was followed by measuring *OD*_600_ at 0 and 24 h. Then, cells were pelleted by centrifugation (20.000 *g*, 15 min, 4°C) and supernatants were stored at −20°C for LC-MS/MS glycomic analysis of HMO depletion profiles.

### Analysis of HMO utilisation profiles using LC-MS/MS

For HMO utilisation profiling, supernatants from the best growing maternal isolates in mYmCFA medium with 1% (w/v) HMOs, based on *OD*_600_ values after 24 h, were analysed by LC-MS/MS as previously described^59^. In short, samples of 40 µL culture supernatant containing 5 µg µL^-1^ HMOs were reduced with 0.5 M NaBH_4_ and 20 mM NaOH at 50 °C overnight and desalted using ZipTip C18 tips (Millipore) packed with a cation exchange resin (AG50WX8, Bio-Rad). After drying (SpeedVac), residual borate was removed by repetitive methanol additions and evaporations. Prior to mass spectrometry, samples were reconstituted in 100 µL milliQ water containing 20 pmol µL^-1^ 3ʹ-sialyl-*N*-acetyllactosamine (Dextra Lab, UK) and analysed by liquid chromatograph-electrospray ionisation tandem mass spectrometry (LC-ESI/MS) using a 10 cm × 250 µm I.D. column, packed with 5 µm porous Hypercarb graphitised carbon particles (Thermo-Hypersil, Runcorn, UK). Glycans were eluted using a linear gradient of 0-80% (v/v) acetonitrile in 10 mM NH_4_HCO_3_ over 46 min at a flow rate of 10 µL min^-1^. The samples were analysed in negative ion mode on an LTQ linear ion trap mass spectrometer (Thermo Electron), with an IonMax standard ESI source equipped with a stainless-steel needle kept at −3.5 kV. Compressed air was used as the nebuliser gas. The heated capillary was kept at 270 °C, and the capillary voltage was −50 kV. A full scan (*m*/*z* 380-2000, two micro scans, maximum 100 ms, target value of 30,000) was performed, followed by data-dependent MS^2^ scans (two micro scans, maximum 100 ms, target value of 10,000) with normalised collision energy of 35%, isolation window of 2.5 units, activation q = 0.25, and activation time 30 ms). The threshold for MS^2^ was set to 300 counts and data were processed using Xcalibur (v 2.0.7). The peak area (the area under the curve, AUC) of each glycan structure was calculated using the Progenesis QI software (Nonlinear Dynamics, Waters Corp., Milford, MA, USA). The AUC of each structure was normalised to the total AUC and expressed as a percentage. The LC-ESI/MS raw data have been deposited in Glycopost under the accession number GPST000504. For the HMO utilisation analysis, a subset of 8 unsubstituted, 9 fucosylated, 5 sialylated HMOs, and 3 putative HMO degradation products were selected, and their relative abundances in individual samples after 24 h of growth, were compared against ten independently analysed HMO controls (non-inoculated mYmCFA supplemented with 1% (w/v) HMOs). Changes in relative abundance profiles of the HMOs were analysed and clustered as a metric for similarities of utilisation and/or enzymatic modification amongst the individual isolates, hierarchical clustering was performed using the “group average” clustering method and Euclidean distance as the distance measure (OriginPro 2023).

### Shotgun metagenomic sequencing, assembly, and annotation

DNA from faeces and enriched cultures was extracted using PureLink Microbiome DNA purification Kit (Fisher Scientific) according to manufacturer’s manual. MGI DNBSEQ sequencing libraries were prepared using the MGIEasy Fast FS DNA library Prep set (MGI Tech Co., Ltd., Shenzhen, China) according to the manufacturer’s protocol. Libraries were sequenced on DNBSEQ-T7 (MGI Tech Co., Ltd., Shenzhen, China) using DNBSEQ-T7RS High-throughput Sequencing Set (FCL PE150) (MGI Tech Co., Ltd., Shenzhen, China) to yield about 40 million paired-end reads per sample. Sample preparation and the whole metagenome shotgun sequencing were performed in Copenhagen. High-performance computing resources were used for all sequencing read analyses described in this section^60^. The quality of sequence reads was assessed using FastQC v0.11.9^61^ and trimmed with Trimmomatic v0.38^62^ (parameters: leading 9, trailing 3, sliding window 4:15, minlen 36). Reads mapped to the human reference genome (GRCh38.p14^8^) were removed using Bowtie2 v2.5.2^63^ for stool samples. Microbial taxonomic profiles were predicted with MetaPhlAn 4 v4.1.0^64^ using the *Bifidobacterium longum* subspecies-resolved database^65^, which converted to GTDB taxonomy r207. Taxa with relative abundance below 0.01% were excluded. Filtered reads were assembled using MEGAHIT v1.2.9^66^ in default mode. Proteins were annotated on the assembled contigs with BAKTA v1.9.3^67^, and carbohydrate-active enzymes (CAZymes)^34^ were predicted with dbCAN v4.1.4^68^.

Filtered reads were aligned to coding sequences (CDS) and contigs using BWA v0.7.17^69^. Mapped reads were sorted and converted to BAM files using SAMtools v1.20. Read counts for proteins were calculated with BEDTools v2.30.0^70^, with gene lengths extracted using SeqKit v2.4.0^71^. These counts were further used to quantify CAZyme family abundances and subfamilies for each sample, using dbCAN-utils script from dbCAN v4.1.4^68^, normalised by genes per million (GPM). Aligned sequences of glycoside hydrolase family 42 (GH42) were extracted from metagenome-assembled genomes (MAGs) retrieved from maternal consortia enriched on HMOs and their multiple specificities including LNT-specific β-galactosidases active on Gal-GlcNAc linkages, β-galactosidases active on various Gal-Gal linkages found in plant-derived saccharides, α-L-arabinopyranosidases were annotated to maximise the accuracy of predictions in the lack of CAZy-defined subfamilies in GH42. A multiple sequence alignment and a phylogenetic tree were generated using MAFFT 7^72^, and the tree was visualised using Interactive Tree of Life v6^73^.

Metagenomic bins were constructed using Vamb v.4.1.3^74^ and SemiBin v.2.1.0^75^, refined with DAS tool v.1.1.6^76^ and dereplicated with CoverM v.0.7.0^77^ at 98% average nucleotide identity. Bins with > 90% completeness and <5% contamination were selected (CheckM2 v.1.0.2^78^). Taxonomic assignment of the bins was performed with GTDB-Tk v2.3.2 (r_220)^79^, and proteins and CAZymes were annotated with BAKTA v1.9.3^67^ and dbCAN v4.1.4^68^, respectively. An overview of the whole metagenome sequencing analysis is shown in Supplementary Fig. 16.

### Microbial diversity analyses

All analyses were performed using R v4.3.2^39^. The Shannon index, calculated using the vegan package v2.6-10^38^, was used as a measure of α-diversity, which reflects the diversity within a single sample by combining richness (the number of species) and evenness (the relative distribution of species). Differences in the Shannon index for both taxonomic diversity and polysaccharide-utilising CAZymes between weaning stages (Pre-weaning, Early weaning, and Late weaning) were compared using a paired Wilcoxon test. Non-metric multidimensional scaling (NMDS) a method for visualising beta-diversity (differences in community composition between samples), was conducted on Bray-Curtis dissimilarities of the relative abundance of microbial species, with selected CAZymes as environmental factors. Species-environment (CAZymes) relationships were assessed using envfit (vegan) with 999 permutations, and significant vectors (*P* < 0.05) were visualised. Needs to include distances calculation between time points.

### Public infant cohort data analyses

Sequence raw data accession IDs for a Swedish infant-mother cohort^1^ was retrieved from the curatedMetagnomicData package v.3.16.1^40^, selecting only samples from exclusively breastfed infants delivered vaginally with no reported antibiotic use. The corresponding raw data were downloaded from Sequence Read Archive (SRA) of the National Center for Biotechnoloyg Information (NCBI), using the NCBI SRA Toolkit v.3.0.0^80^, and processed and analysed using the same pipeline as applied to our Milkome cohort (Supplementary Fig. 16), as described above.

### Search for HMO utilising genes in Human Gut Microbiome (HGM) Catalogue

To investigate the prevalence of HMO utilising genes in bacteria of the GM, we identified glycoside hydrolase family 112 (GH112) and GH136-like proteins within the HGM protein catalogue, consisting of 170,602,708 proteins clustered at 100% amino acid similarity (version 2.0, downloaded in December 2021) from 286,997 genomes^44^. Hidden Markov model (HMM) profiles for GH112 and GH136, sourced from dbCAN-HMMdb-V11 (downloaded in August 2022)^68^, were used to identify GH112- and GH136-like proteins via the Hidden Markov Model (HMM) search function from HMMER with default settings^81^. All matching sequences were then retrieved, and their coverage relative to the query profiles was calculated. Curation threshold was set as E-value cutoff ≤0.001, coverage thresholds of ≥80% (GH112) and ≥60% (GH136), and minimum size of 600 and 300 amino acids, respectively. A multiple sequence alignment and a phylogenetic tree were generated using MAFFT 7^72^, and the tree was visualised using Interactive Tree of Life v6^73^. Active site conservations of retrieved protein sequences were mapped on the identified active sites of the *Bifidobacterium longum* JCM 1217 (GH112, PDB ID: 2zuv[A],)^82^ and *Eubacterium ramulus* ATCC 29099 (GH136, PDB ID: 6KQS[A]) ^36^ and illustrated with WebLogo 3^83^. The abundance of GH112- and/or GH136-encoding genes was calculated by mapping the representative proteins from the catalogue to their corresponding clustered proteins and their genomic origins using a provided mapping file^44^.

### Statistical analyses and visualisation

All statistical analyses and visualisation were performed using R v4.3.2^39^. The growth of the first enrichment, conducted individually for each donor, was compared to no-carbohydrate-added controls and carbohydrate-supplemented cultures, with data pooled across different weaning stages and maternal samples. Data were pooled across donors at each time point and analysed using generalised linear mixed models implemented in glmmTMB v1.1.10^84^, with random effects accounting for donor identity, substrate supplementation of the medium, and duplicates at each time point. For enrichment II, growth was compared to no-carbohydrate-added controls and to carbohydrate-supplemented cultures, analysing data per time point for each donor individually using two-tailed t-tests with unequal variances. The number of CAZymes per million, categorised by carbohydrate type, was compared using the Wilcoxon rank-sum test in R v4.3.2^39^ and visualised with the ggpubr package v0.6.0^85^. To assess whether the median values of fibre utilisation genes, normalised by gene per million, were greater than zero in exclusively breastfed infants from Swedish cohort^1^, *P*-values were calculated using a one-sided Wilcoxon signed-rank test^39^. Differences in community composition across timepoints within infant faecal samples were evaluated using PERMANOVA using adonis2 from vegan package^38^ based on Bray-Curtis dissimilarity with 9,999 permutations. Additionally, group-specific differences in faecal and enriched consortia from mothers and infants were assessed via PERMANOVA^38^, with each age and substrate group compared against all others. Plots were generated with ggplot2 v3.5.2^86^ and heatmaps by pheatmap v.1.0.12^87^. Carbohydrate utilisation profiles of high quality MAGs were visualised using ComplexUpset v1.3.3^88^

## Data availability

The sequencing data are publicly accessible in the European Nucleotide Archive under project ID XXXX. The glycomics MS raw files have been deposited in the GlycoPOST database under the ID GPST000504

## Code availability

Codes used for the study are available at https://github.com/yunso123/MILKOME.

## Acknowledgements

The study was funded by a Research Fund Denmark, Natural Sciences (FNU Project 2 grant no. 1026-00386B) to M.A.H. C.J was partially employed on a Novo Nordic Foundation Interdisciplinary Synergy Programme (grant no. NNF22OC0077684). We express our deep gratitude to all the infants and mothers for their participation in this study. We are grateful to the DTU Computing Center for providing high-performance computing resources. We thank Maria Louise Leth for her contribution in translating the Milkome cohort protocol to Danish and the undergraduate students Sofus Boisen, Nicoline Bølling Bjørnskov Jensen, Barbara Borcic, and Lucija Backov for their work within the project. We want to express our gratitude to Associate. Prof. Manimozhiyan Arumugam and Dr. Camila Alvarez-Silva for their kind help at the early stage of the project. We also would like to thank to Senior Researcher Dr. Jesper Holck for help with the initial HMOs glycomic analysis. Figures 1 and 3, Supplementary Figures 1,2, and 6 were partially created in BioRender (So, Y. (2025) https://BioRender.com/6clwofm).

## Author contributions

M.A.H., Y.S. and M.J.P conceptualised this work. L.A. and S.S.K. were responsible for the cohort recruitment. Y.S. and M.J.P developed methodology on the microbial work and M.J.P. has developed the HMO preparation. C.J. and S.T. provided methodology and infrastructure for the glycomic analyses, while K.K provided methodology and infrastructure for the metagenomic shotgun sequencing. Y.S. and M.J.P conducted the investigations. Y.S. carried out the analyses with the help of M.J.P., C.E., I.C, and S.B. Y.S. created the visualisations with the help of M.J.P, who performed the mother isolate HMO analysis and visualisation. M.A.H acquired funding and carried out the project administration. M.A.H. and Y.S. wrote the first draft of the manuscript with input from C.J. and M.J.P., who wrote the sections related to HMO utilisation by mother. All authors contributed to the manuscript editing and approved the final manuscript version.

## Additional information

The Supplementary Information is available online

## Ethical considerations

Volunteers were enrolled based on informed consent and the study was approved by the ethical committee of the Capital Region under the approval of the Capital Region ethical committee (Study registration ID: H-21067700 and registered in Clinicaltrials.gov under accession ID NCT07026526)

## Supplementary Information Inventory

### Supplementary Results

#### The HMO utilisation apparatus in the GM

Lacto-*N*-biosidases of GH136 break down the abundant lacto-*N*-tetraose (LNT) backbone or its fucosylated versions in type I HMOs (Supplementary Figs. 1d and 2). Lacto-*N*-biosidases of GH136 cleave LNT into lacto-*N*-biose (LNB, Galβ1,3GlcNAc) and lactose. The second specificity in GH136 enzymes mediates the cleavage of fucosylated LNB forms (Lewis a/b antigens) from α1,4-fuocylated LNT, *e.g.*, lacto-*N*-fucopentaose II (LNFP II) or the double fucosylated form. (Supplementary Figs. 1 and 2). The most common route to catabolise the LNB product, released in tandem of GH136 activity, is via LNB phosphorylase (GH112). The degradation of LNT can also be initiated by distinct β-galactosidases (GH42), *e.g.*, from *B. infantis* that cleave the non-reducing galactosyl unit (Supplementary Fig. 2). Of note, β-galactosidases/α-L-arabinopyranosidases specific for plant-derived substrates^3^ populate different clades of the GH42 phylogenetic tree (Supplementary Fig. 9c), justifying the inclusion of this family in our comparative analysis. Several other families mediate HMO degradation but also harbour overlapping activities on host-derived mucin *O*-glycans (and/or related *N*- and *O*-glycoconjugates), including α-1,3/4/6-fucosidases (GH29), α-1,2-fucosidases (GH95), sialidases (GH33) and *N*-acetylhexosaminidases (GH20). Finally Lacto-*N*-biosidase^4^ of GH20 degrade LNT, similarly to GH136, resulting in LNB and lactose (Supplementary Fig. 2).

#### Medium optimisation for the culturomics analyses

We initially tested the yeast extract-casitone, which is a casein hydrolysate-fatty Acids medium (YCFA) that supports broad growth of GM taxa^5^, which was supplemented with low amount of porcine gastric mucin (0.05% w/v) to provide an adhesion surface to mucus-associated GM. To suppress baseline growth on the medium, we decreased yeast extract, utilised by certain *Bacteroides*^6^, by 2.5-fold (mYCFA). This medium was used for the initial enrichment (enrichment I) of the faecal consortia after supplementation with HMOs, mucin, xyloglucan (XG), or 1:1 blends of arabinoxylan (AX) and pectin (PEC), or galactomannan (GMN) and gum Arabic (GA), each at 0.5 % (w/v) (Supplementary Tables 4-6). A mixture of 0.5 % (w/v) of each glucose, galactose, and fucose was additionally used for broad recovery of GM, irrespective of complex carbohydrate utilisation ability. A fibre blend was used to minimise handling time of the fresh faecal consortia for best microbial recovery in the first enrichment (Enrichment I) (Supplementary Fig. 5, Supplementary Table 6). No inoculum and no carbohydrate supplementation cultures served as contamination and baseline (growth on medium) controls. Growth was monitored by changes in *OD*_600_ and pH value (*Δ*pH), resulting in similar trends (Supplementary Fig. 5). Baseline growth in the no carbohydrate controls was still observed, likely attributed to protein fermentation. Thus, the content of Casitone (protein source) was reduced by 50% (mYmCA) as compared to mYCFA (Supplementary Table 4). The optimised medium mYmCFA showed lower baseline growth and was used to repeat the enrichments (enrichment II, Fig. 2a) to minimise the propagation of taxonomic groups that grow on the medium, but not on the supplemented carbohydrates. This was crucial to reduce false positives in the tandem metagenomic analyses of the enriched consortia.

#### HMO utilisation by cohort mother HMO utilising isolates

We started by isolating 50 colonies from each mother (n=7), except for mothers C (n=38 strains), E and M (n=35 for both), where a lower number of isolates was observed on the plate. The isolates were re-streaked twice on the HMO-agar plates, attempting to obtain single strain isolates. These isolates were regrown on the same medium in 96-well plates, with 188 isolates showing significant growth (*P*-value < 0.01) as compared to the no-carbohydrate supplemented medium (NC). Of all isolates, 19 % (n=36) grew poorly (*OD*_600_= 0.2-0.3), while 93 exhibited moderate to high growth (*OD*_600_ > 0.3), all compared to NC controls (Supplementary Figs. 11 and 12a). Growth levels and distribution of individual isolates were similar across donors, except for one mother (D), exhibiting significantly lower growth median), underscoring HMO utilisation as a prevalent trait amongst the maternal GM (Supplementary Fig. 12b).

Then, we selected isolates with growth levels *OD*_600_= 0.1–0.79, relative to no-carbohydrate controls. The isolates were from mothers A (n= 28), B (n= 27), C (n= 29), D (n= 1), E (n= 29), and G (n=23). We supplemented HMOs purified from mothers’ milk in the standard growth medium, then performed an LC-MS analysis to annotate 68 top abundant HMO structures. Based on this, we selected 22 representative HMO structures (Supplementary Fig. 13) and analysed changes in the HMO profiles in spent culture media after growth of the HMO-utilising isolated, as compared to the non-inoculated controls (n=10) as a proxy for HMO utilisation patterns. The HMOs for this glycomic analysis were chosen based on abundance, backbone chain type, and patterns of fucosylation/sialylation. Following an initial screen, three potential HMO degradation products, that were detected in the spent culture supernatants but not in the non-inoculated medium, were also included in the analysis (Supplementary Fig. 13).

Based on the differential HMO profiles, the maternal isolates were clustered into five groups (Supplementary Fig. 12c). Group I showed efficient degradation of undecorated HMO backbones (LNT/LN*n*T, lacto-*N*-*neo*tetraose), including those branched with β1,6-*N*-acetylglucosamine unit (Fig. 7 and Supplementary Fig. 1). Both group II and III exhibited α-fucosidase activity but were distinguished by distinct substrate preferences (Fig. 7 and Supplementary Fig. 12c). Group II members could remove fucose from both α1,2- and α1,3/4-linkages in simple structures such as fucosyllactose (FL) and more complex structures, including fucosylated, branched LNT/LN*n*T backbones. By comparison, group III members exhibited α-fucosidase activities that targeted terminal α1,2-linkages in abundant 2ʹ-FL and α1,3-fucosyl units in larger, complex structures, but notably lacked activity against 3-FL (Fig. 7). Members of both groups displayed limited capacity to utilise LNT/LN*n*T structures. However, group II members exhibited a modest ability to access branched LNT/LN*n*T structures following defucosylation, likely facilitated by a putative β-galactosidase activity on β1,4-linkages in *N*-acetyllactosamine, referred to as type II chains (Supplementary Fig. 1e). This trend also applies to groups IV and V. Group IV exhibited weaker fucosidase (targeting α1,2- and α1,3/4-linked fucose caps) and sialidase (targeting α2,3- and/or α2,6-linked sialic acid caps) activities compared to groups II and III and were unable to directly access LNT/LN*n*T, while some members appeared to utilise galactosyl-lactose-extended HMOs (Supplementary Fig. 8), likely via an extracellular β-galactosidase activity. In contrast, group V isolates, lack both fucosidase and sialidase activities and were similarly unable to directly access LNT/LN*n*T. However, these isolates were able to utilise larger, extended, or branched LNT/LN*n*T structures without terminal decorations, resulting the second-highest median growth (Fig. 7). This pattern may likely emerge from extracellular β-galactosidase and β-*N*-acetylglucosaminidase activities and subsequent uptake of the released monosaccharides. These results reiterate the importance of the enzymatic and uptake machinery of targeting abundant LNT/LN*n*T backbone structures, as compared to exo-glycosidase activities that cleave fucose/sialic acid caps of HMOs. Thus, group I was significantly overrepresented, while groups II and III were significantly underrepresented among all isolates (Supplementary Fig. 12d) or when examining the 20 best-growing isolates (Supplementary Fig. 12e).

#### Clustering and active site analysis of HMO-active GH112 and GH136 enzymes

After analysing the taxonomic distribution of the HMO-active GH112 and GH136 sequences, retrieved from the human gut microbiome catalogue^2^, we analysed their catalytic apparatus. To this end, we performed multiple sequence alignments together with characterised GH112 and GH136 sequences from CAZy (www.CAZy.org^4^) as references and generated a phylogenetic tree of each family (Supplementary Fig. 15a-d). Most identified GH112 genes clustered with LNB/GNB (galacto-*N*-biose) phosphorylases. Similarly, the GH136 enzymes clustered with lacto-*N*-biosidases, whereas a large clade was associated with the cleavage of fucosylated LNT chains, *e.g.*, in LNFP II, lacto-*N*-difucohexaose I (LNDFH I), and lacto-*N*-difucohexaose II (LNDFH II) (Supplementary Fig. 1 and 15c,d). Sequence logos revealed high conservation of substrate-binding and catalytic residues, in line with the predicted function for these sequence (Supplementary Fig. 15e,f). Of note, GH136 members are heavily under-characterised, and additional specificities may be harboured within this family.

### Supplementary Figures

**Supplementary Fig. 1.**
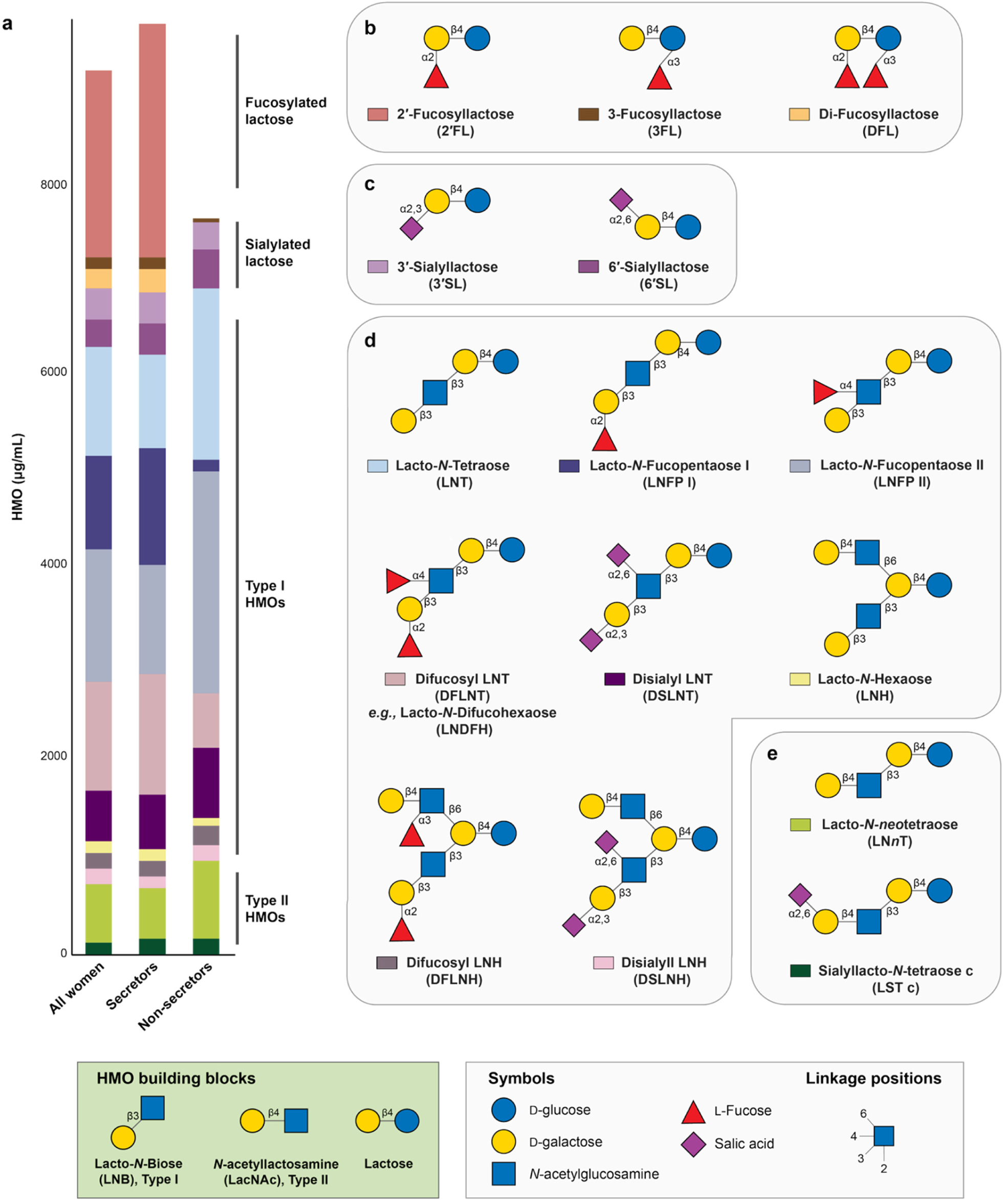
Human milk oligosaccharide distribution and structures. **a** HMO absolute abundances calculated from a previously published study from mothers (n =410), including secretors (n=316) and non-secretors (n=94). Samples were collected between two-weeks and five-month postpartum. The values represent averaged data from donors across eight regions: Ethiopia, Gambia, Ghana, Kenya, Peru, Spain, Sweden, and the US. Data were sourced from McGuire *et al* (ref. 89 in the main article) and calculated by the author. **b** Fucosyllactoses **c** Sialyllactoses **d** Examples of Type I HMOs. **e** Examples of type II HMOs. Structures are shown according to the Symbol Nomenclature for Glycan (SNFG).

**Supplementary Fig. 2.**
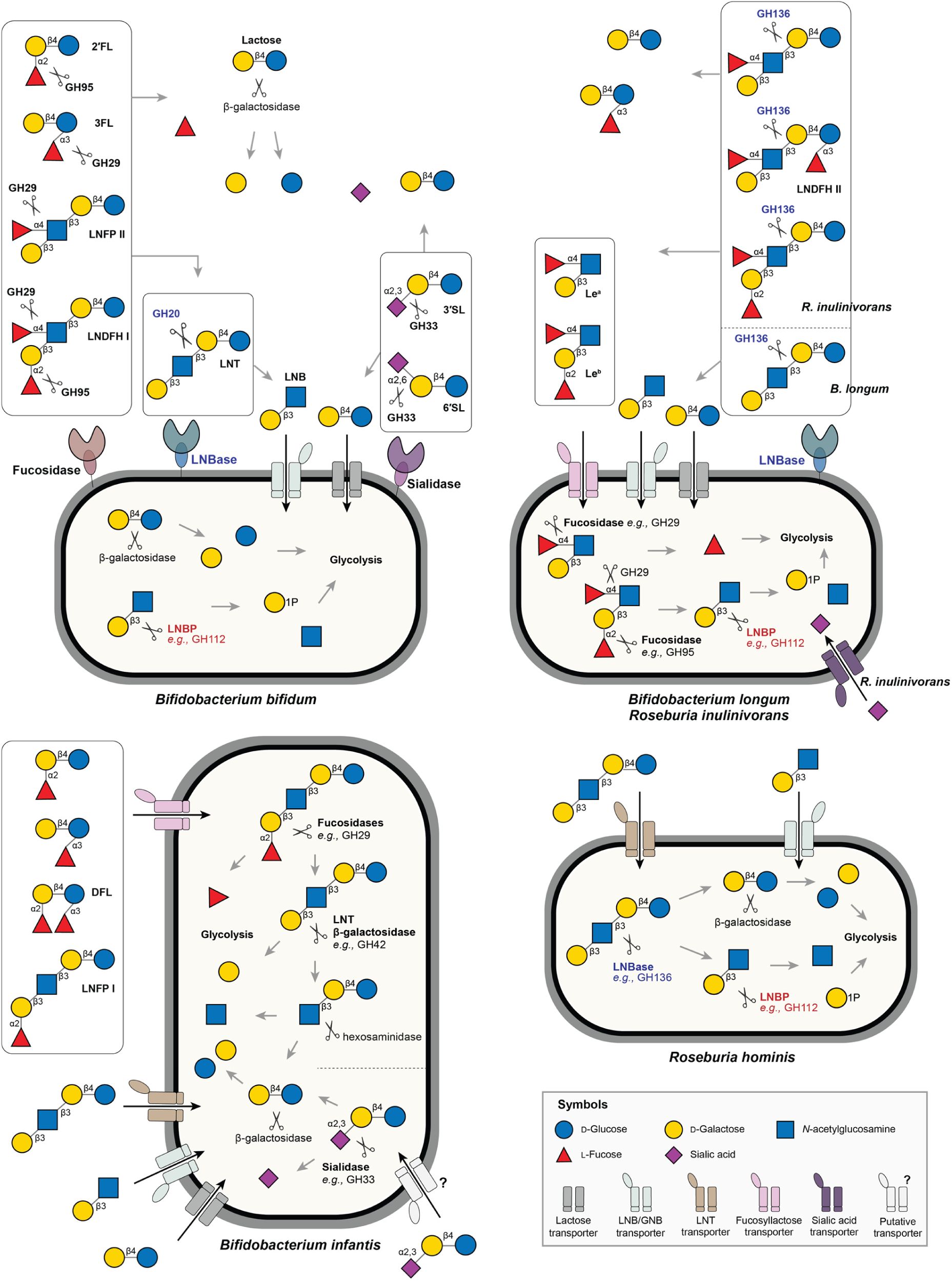
Schematic HMO utilisation models based on described GM members. **a** Mainly extracellular utilisation systems of *B. bifidum*. **b** Systems for the utilisation of LNT by *B. longum* and fucosylated LNT forms by *R. inulinivorans* and related Clostridiales. **c** and **d** Intracellular systems of *B. infantis* and *R. hominis*, respectively. The references that reported these pathways are cited in the main manuscript text.

**Supplementary Fig. 3.**
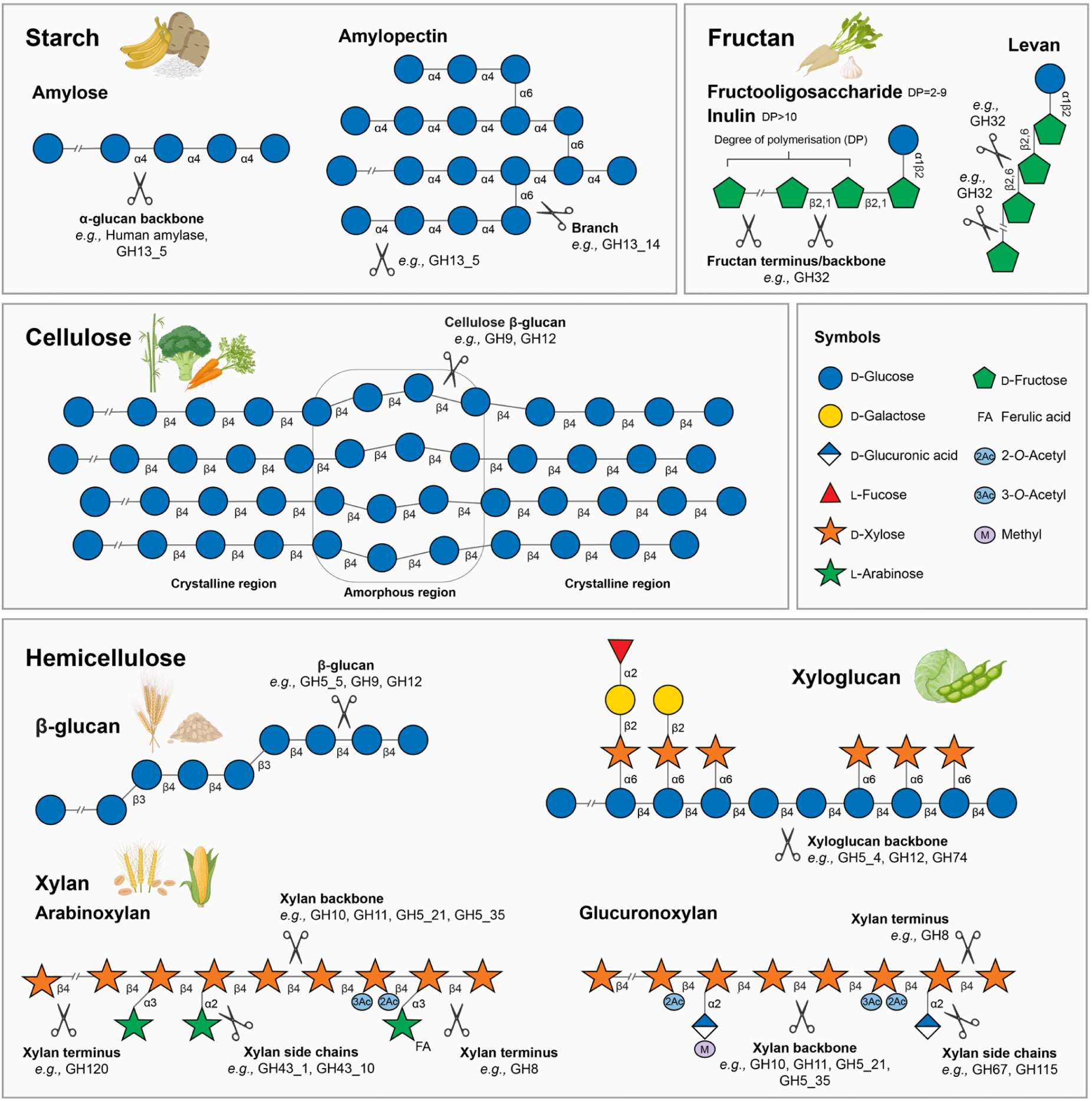
Plant polysaccharide degradation by selected CAZymes. Schematic illustration of plant energy storage and structural polysaccharides relevant to human diet. Starch, composed of amylose and amylopectin, is the primary energy reserve in plants and is the prime calorie source in human diet *e.g.*, cereals, potatoes, and bananas^7^. Starch can be either accessible or resistant to human digestion. Resistant starch transits to the colon and acts as a dietary fibre. Fructans, serving a similar storage role in some plants, are represented by fructooligosaccharides and inulin, which are common in chicory root, onions, and garlic, while levan is found in fermented soybean (Natto)^8^. Structural polysaccharides include cellulose and the hemicelluloses β-glucan^9^ (from oats and barley), xyloglucan^10^ (from legumes, fruits, and vegetables), two type of xylans (from wheat and corn); arabinoxylan and glucuronoxylan^11^. Selected CAZymes specific to each plant-derived polysaccharide are depicted based on curated data from www.CAZy.org^4^. Structures are shown according to the Symbol Nomenclature for Glycan (SNFG)^12^.

**Supplementary Fig. 4.**
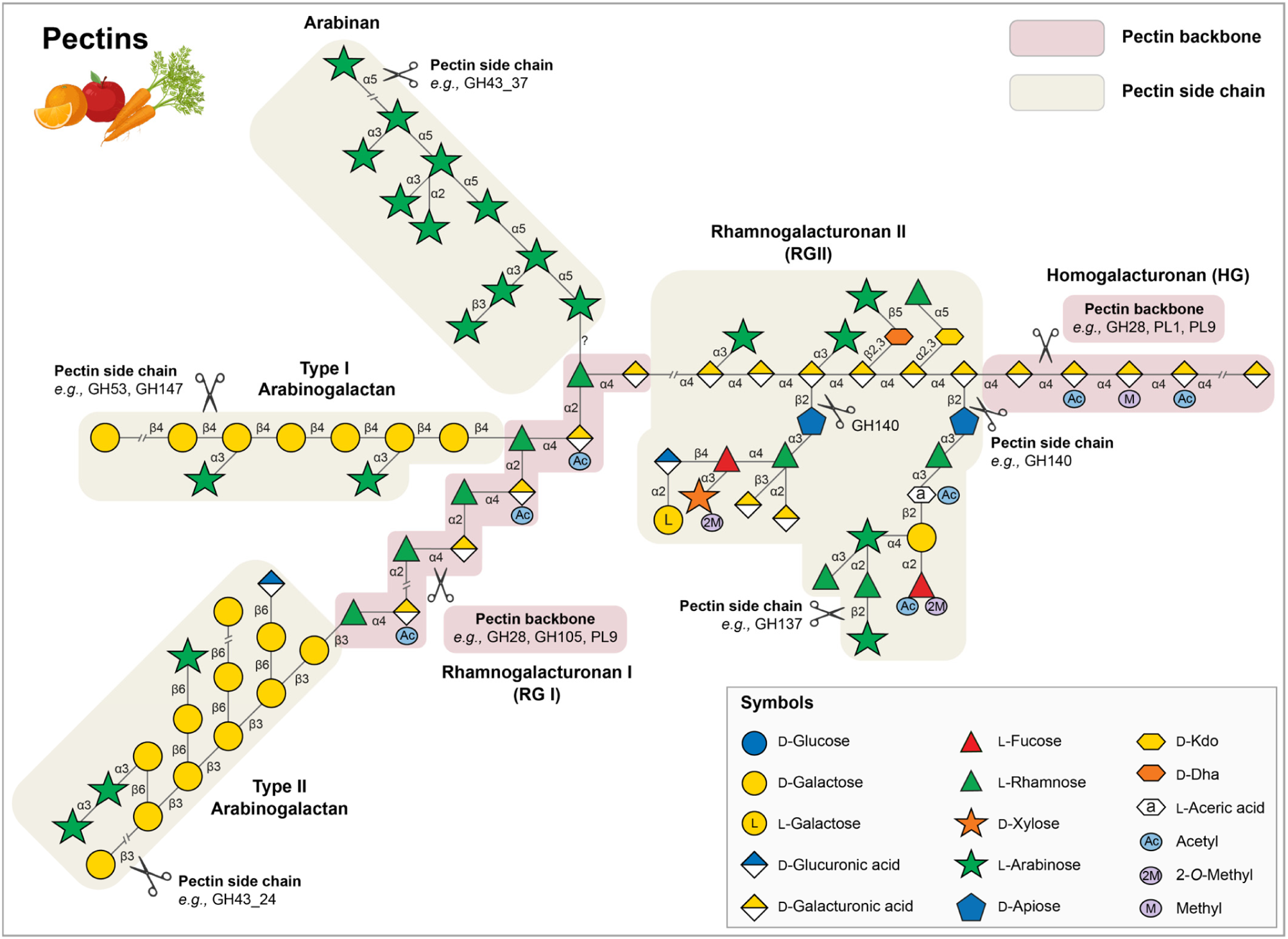
Pectin degradation by selected CAZymes. Pectin consists mainly of two types of backbone structures: homogalacturonan (HG), rhamnogalacturonan (RG) I. The HG backbone can be further substituted by RG II, while the RG I backbone can be decorated by sidechains like arabinogalactan and arabinan^13–16^. Additional xylo-galacturonan domains, where the HG backbone is decorated with xylosyl residues is not included in this simplified schematic representation of the main pectin structures. Schematic structures of pectin backbones and sidechains are shown according to the Symbol Nomenclature for Glycan (SNFG)^12^.CAZymes that are active on pectin backbones and side chains were selected based on curated data from www.CAZy.org^4^.

**Supplementary Fig. 5.**
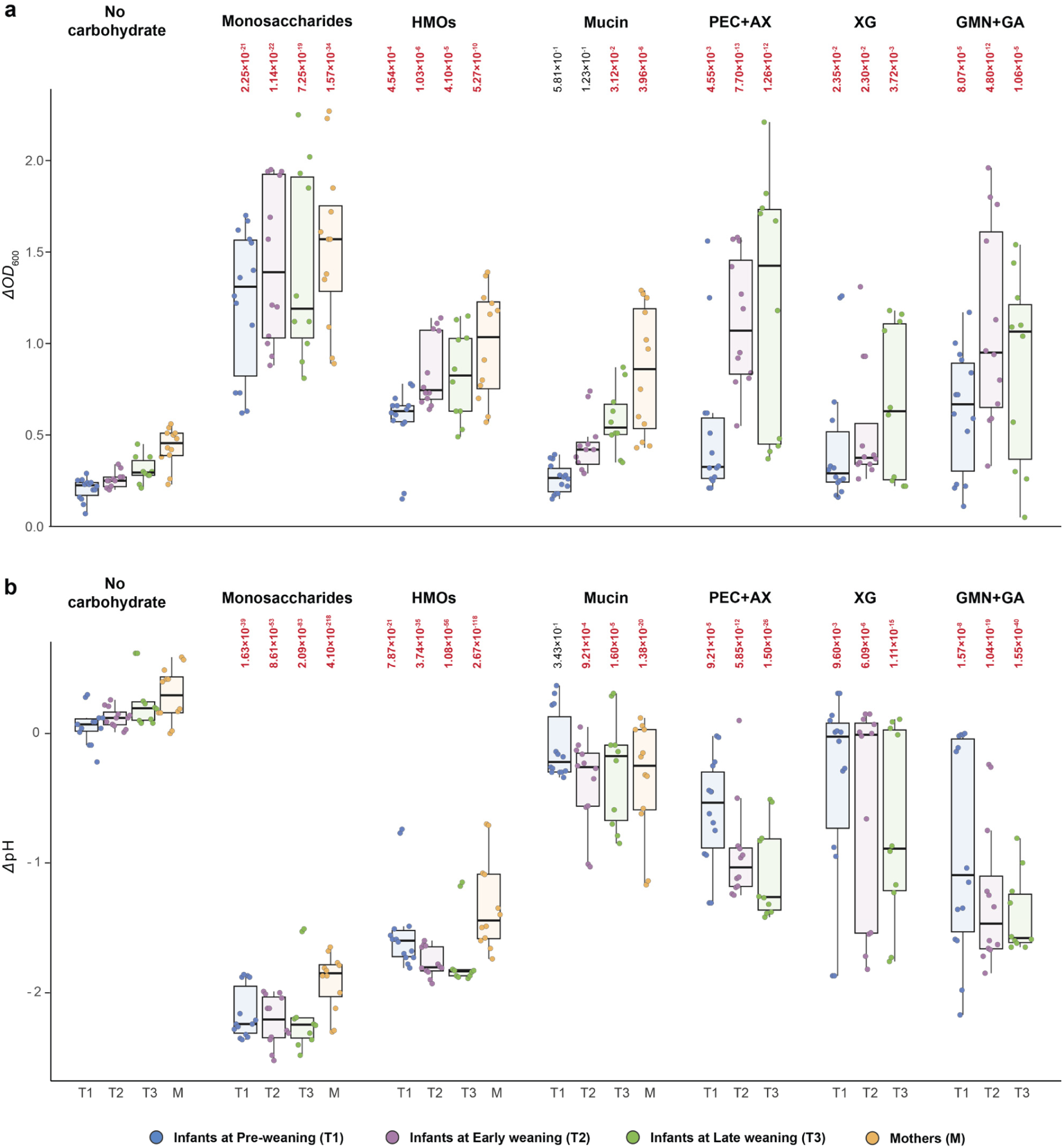
Enrichment of faecal consortia of on carbohydrate substrates. **a** The *OD*_600_ changes from 0 h to 24 h are indicated after enrichment of infant faecal samples at three weaning stages on carbohydrate substrates; No-carbohydrate source added negative controls (NC), monosaccharides (0.5% w/v of each glucose, galactose, fucose), 1 % (w/v) HMOs, 0.5% (w/v) mucin, 0.5% (w/v) pectin (PEC) + 0.5% (w/v) arabinoxylan (AX), and 0.5% (w/v) xyloglucan (XG). **b** The pH changes from 0 h to 24 h. The data are presented as whisker plots and p-values as compared to NC were calculated using linear mixed models implemented in glmmTMB v1.1.10 and the *P*-values showing the statistical significance relative to the controls are shown on top of each substrate.

**Supplementary Fig. 6.**
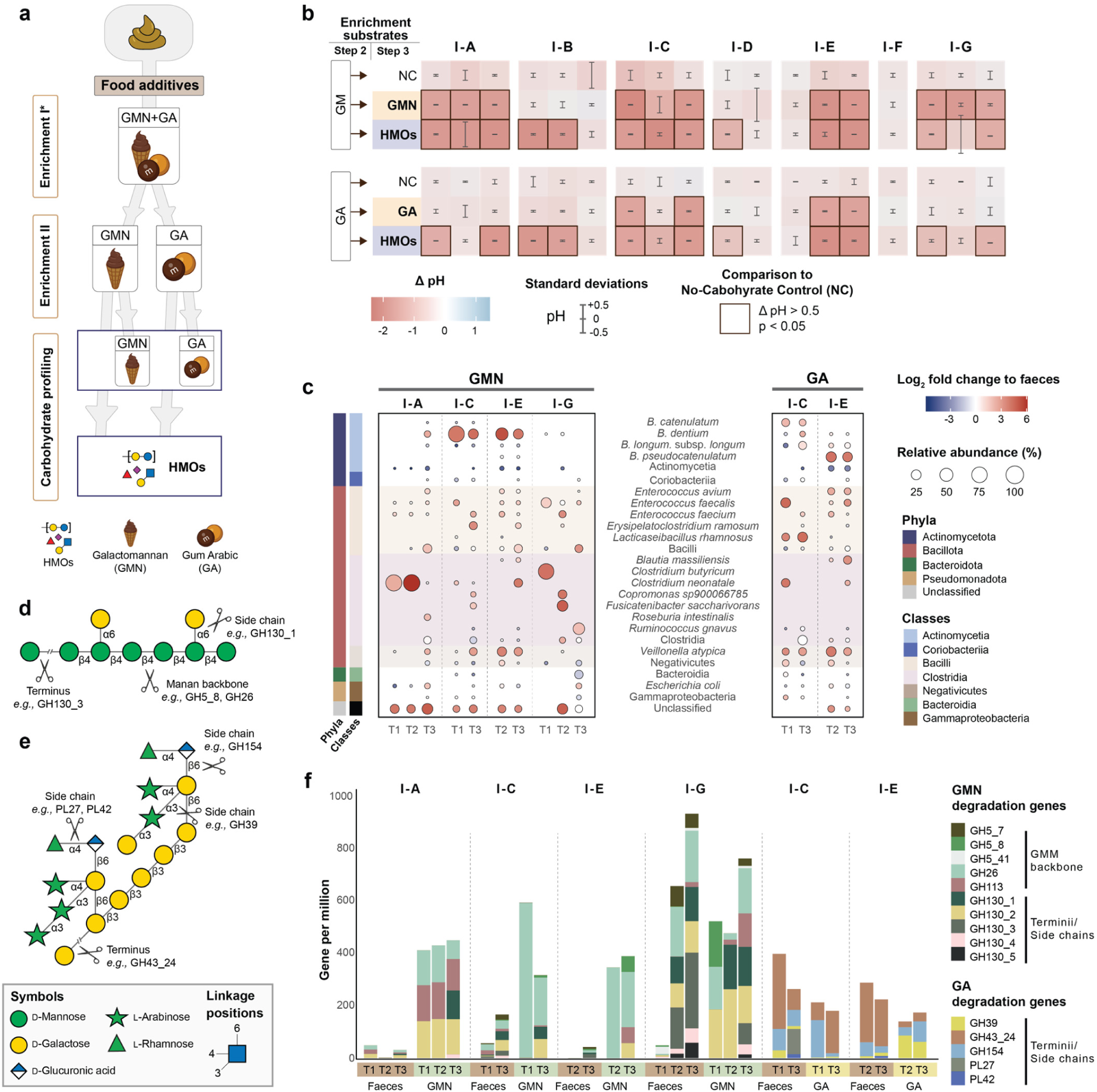
Enrichment of faecal consortia on galactomannan and gum Arabic. **a.** Workflow for faecal sample enrichment on food additives, galactomannan (GMN) and gum Arabic (GA). Enrichment I was performed on a yeast extract-reduced YCFA medium (mYCFA), whereas enrichment II was performed on an optimised medium (mYmCFA), with reduced yeast extract and protein (casitone). **b** Changes of pH after a 24-hour enrichment on GMN and GA. The enriched consortia on the food additives were re-grown on HMOs to assert growth on both HMOs and dietary fibres. The *P*-values were calculated using t-test with the vegan package. **c** Taxonomic composition of the enriched consortia. **d** Schematic representation of galactomannan structure. **e** Schematic representation of gum Arabic structure. **f** The normalised abundance (gene per million) of GMN and GA degradation genes in faeces samples, GMN, and GA enriched consortia. References to the used software in the main article.

**Supplementary Fig. 7.**
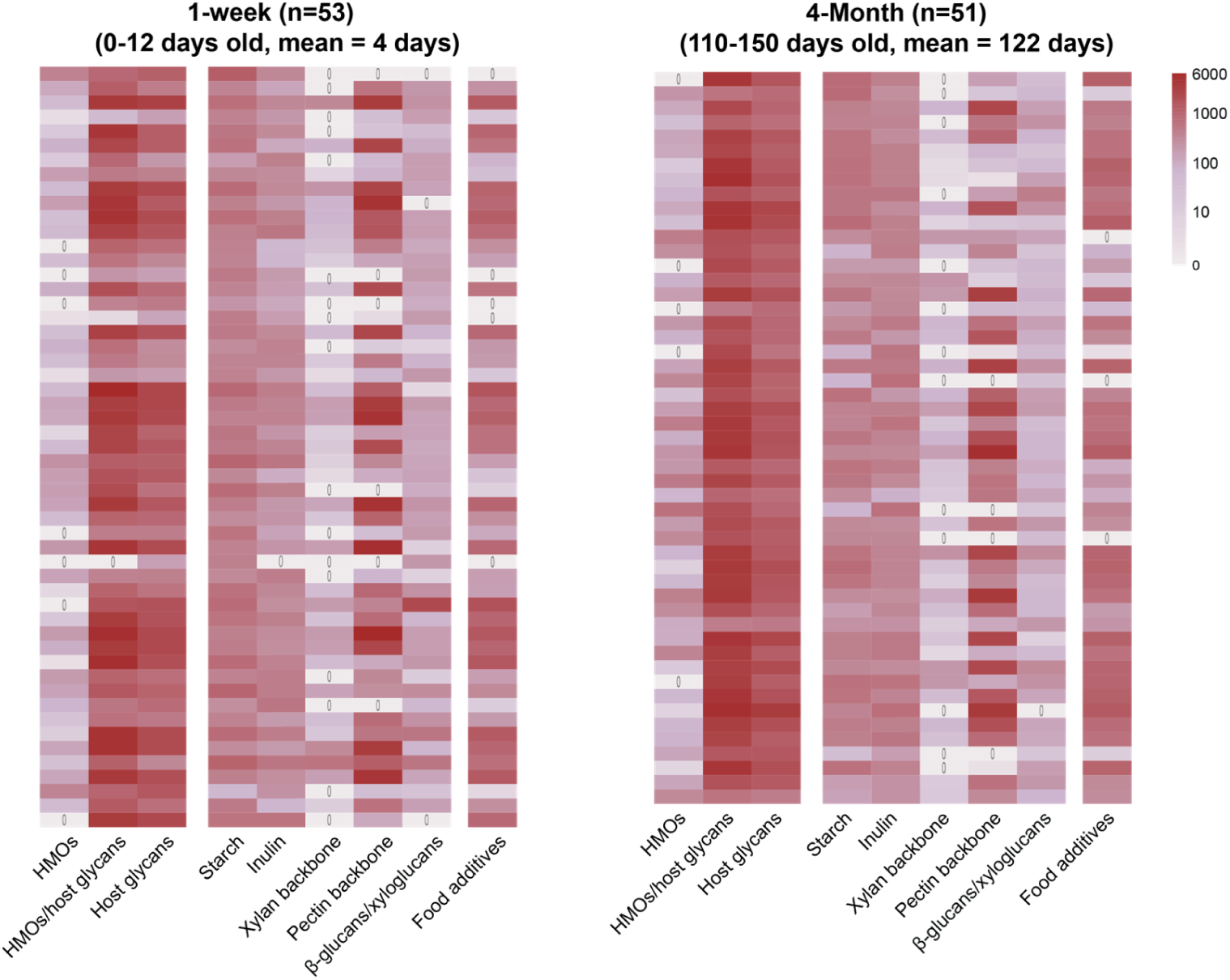
Fibre utilisation genes per infant in exclusively breastfed infants. The data is computed from a Swedish early life cohort (n=98)^1^. Exclusively breastfed infants, delivered vaginally with no reported antibiotic use, at one week (n=53, 0-12 days old, mean=4 days) and 4 months (n=51, 110-150 days old, mean=122 days) are selected and used for the analyses. The number of genes encoding CAZymes that mediate the utilisation of each complex carbohydrate substrate was identified in exclusively breastfed infants, delivered vaginally with no reported antibiotic use, during the first week of life (n=53) and at four months (n=51). Each row in the dataset represents an individual. The number of selected CAZyme families per substrate is: HMOs (n=2), HMOs/host glycans (n=9), host glycans (n=26), starch (n=14), inulin (n=1), xylan backbone (n=4), pectin backbone (n=16), β-glucans/xyloglucans (n=15), and food additives (n=20). The selected CAZyme families for each substrate are provided in Supplementary Tables 1 and

**Supplementary Fig. 8.**
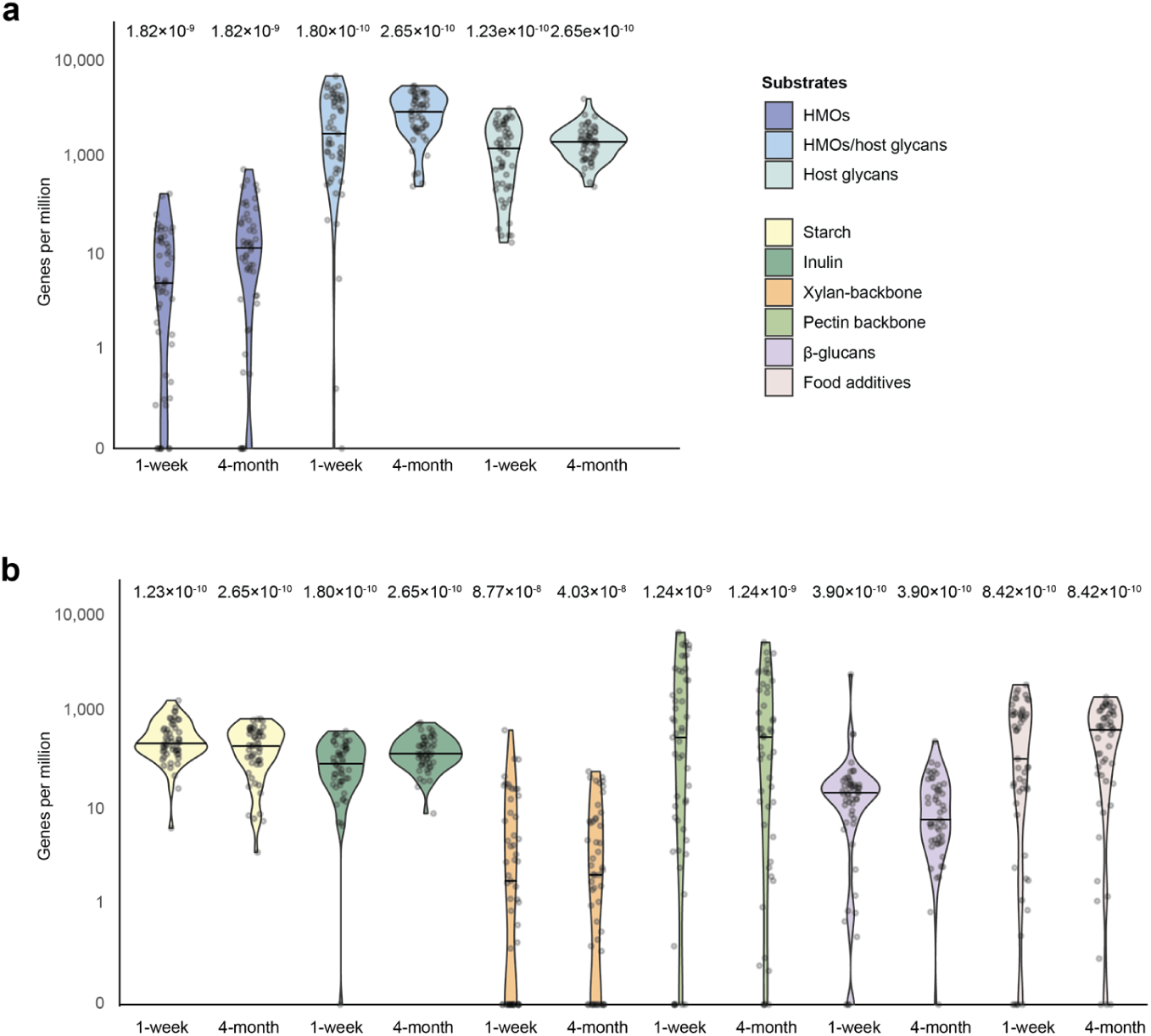
Fibre utilisation genes per fibre type in exclusively breastfed infants. The data is computed from a Swedish early life cohort (n=98)^1^. Exclusively breastfed infants, delivered vaginally with no reported antibiotic use, at one week (n=53, 0-12 days old, mean=4 days) and 4 months (n=51, 110-150 days old, mean=122 days) are selected and used for the analyses. The number of genes encoding enzymes mediating utilisation of each complex carbohydrate, identified in the exclusively breastfed infants in the first week of life and 4 months. **a** HMOs (n=2), HMOs/host glycans (n=9), host glycans (n=26) are shown. **b** Plant derived carbohydrate utilisation of exclusively breastfed infants is shown: starch (n=14), inulin (n=1), xylan backbone (n=4), pectin backbone (n=16), β-glucans/xyloglucans (n=15) and food additives (Galactomannan and gum Arabic, n=20). See Supplementary Tables 2 and 3 for comprehensive details on CAZyme families associated with the degradation of each carbohydrate. *P*-values were calculated using a one-sided Wilcoxon signed-rank test to assess whether median values were greater than zero.

**Supplementary Fig. 9.**
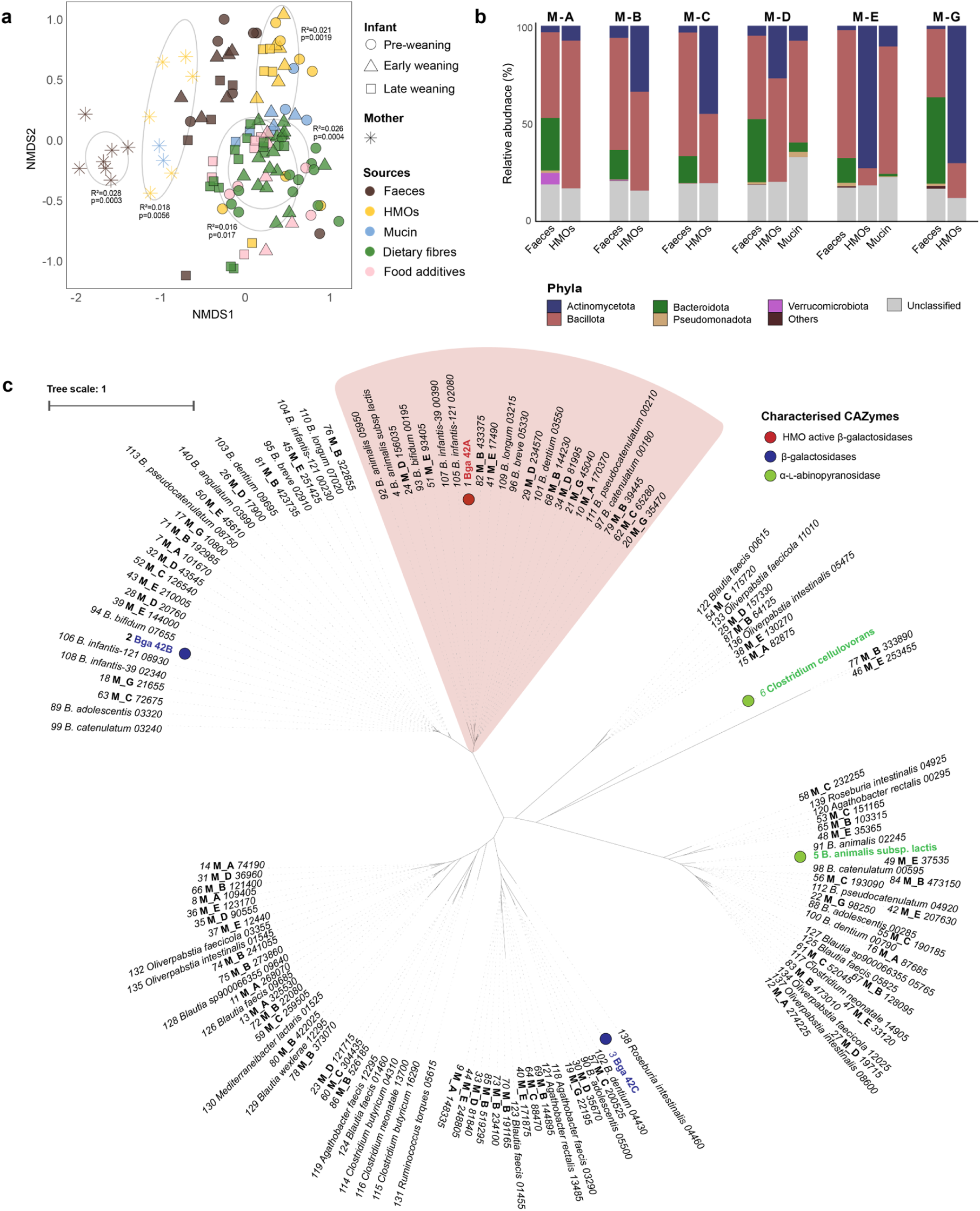
Enrichment of mother faecal consortia on HMOs and mucin. **a** Non-metric multidimensional scaling (NMDS) plot showing community similarity at the species level between mother and infant faeces and enriched consortia, showing the segregation of both the mother faecal communities and enriched cultures from infant samples. Group-specific significance was assessed using PERMANOVA (adonis2, Bray-Curtis dissimilarity, 9,999 permutations), which tests whether overall community composition differs between groups. Each age and substrate group was compared against all others. Only groups with *P* < 0.05 are shown with ellipses and labelled with R² and *P*-values. **b** Phylum level composition of the faecal samples and the enriched consortia. **c** Phylogenetic tree of GH42 enzymes from mother faecal communities enriched on HMOs and MAGs from both faeces as well as enriched consortia. Characterised GH42 enzymes are included as a reference.

**Supplementary Fig. 10.**
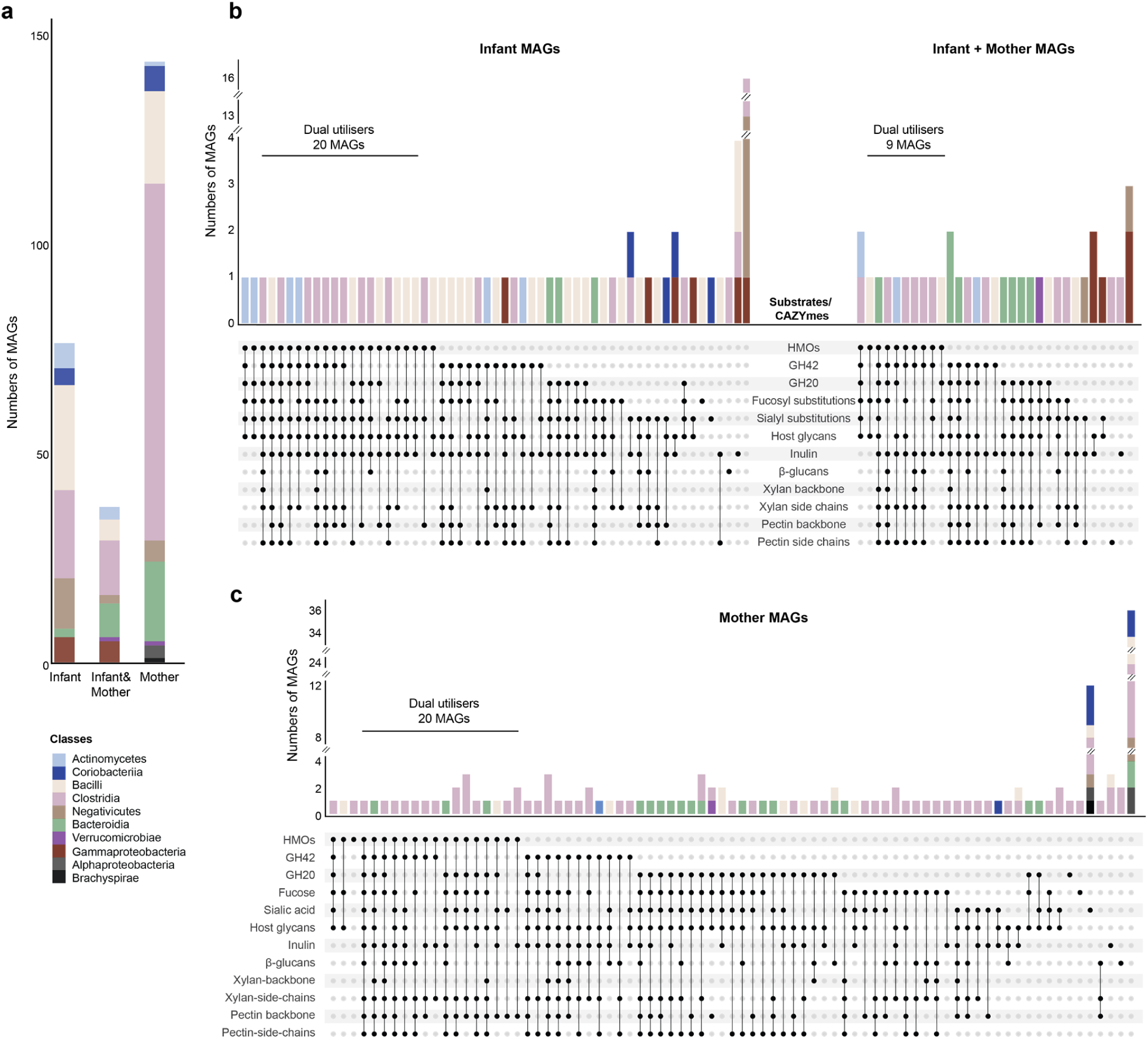
Cohort MAGs and their predicted carbohydrate utilisation profiles. **a** Numbers of MAGs retrieved from infants, mothers or shared between infants and mother. **b** Predicted carbohydrate utilisation potential of MAGs from infants and those shared between infants and mothers. The classification as HMO utilising is based on the presence of GH112 and/or GH136. However, the lack of these enzymes does not exclude HMO utilisation via other routes, *e.g.* GH20 enzymes or combinations of fucosidases/sialidase/β-*N*-acetylhexosaminidases/β-galactosidases (see Supplementary Fig. 2 for details). Therefore, the HMO-utilisers and dual utilisers are likely underestimated. According to these criteria, 20 infant MAGs were identified as dual utilisers (HMOs + dietary fibres), and 9 dual utilisers were from MAGs from both infants and mothers. **c** Predicted carbohydrate utilisation potential of MAGs found exclusively in mothers, including 20 MAGs predicted to be dual utilisers. The number of selected CAZyme families per substrate is: HMOs (n=2), fucosyl substitutions (n=3), sialyl substitutions (n=5), host glycans (n=26), starch (n=14), inulin (n=1), xylan backbone (n=4), pectin backbone (n=16), β-glucans/xyloglucans (n=15), and food additives (n=20). The selected CAZyme families for each substrate are provided in Supplementary Tables 1 and 2.

**Supplementary Fig. 11.**
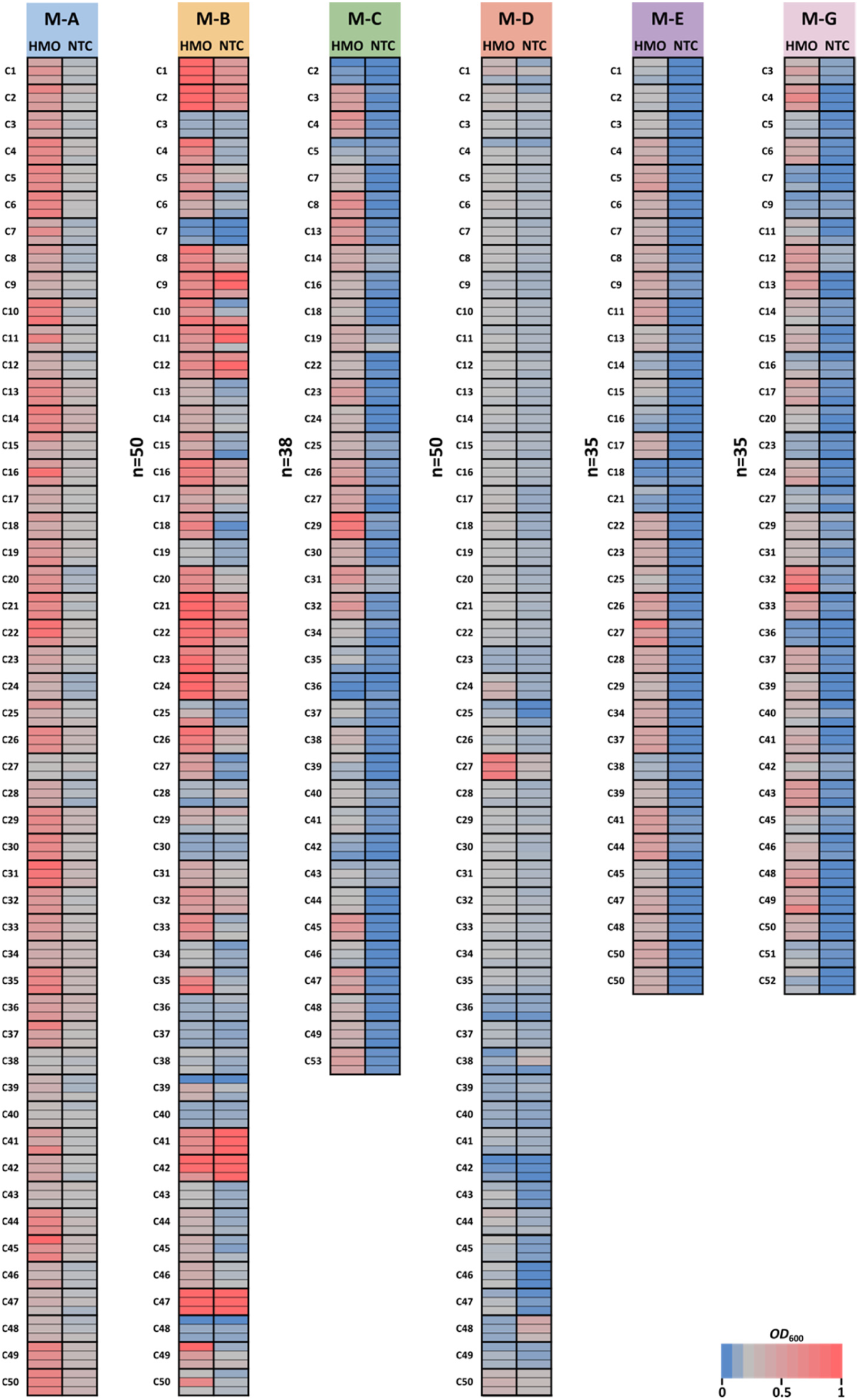
Growth of bacterial maternal isolates on HMOs. Growth of bacterial isolates from mothers A, B, C, D, E and G on mYmCFA supplemented with 1 % (w/v) HMOs after 24 h including a no-carbohydrate control (NTC). Growth experiments were performed in three independent biological replicates (n=3). Source data are provided as a Source Data file labelled with the corresponding figure number.

**Supplementary Fig. 12.**
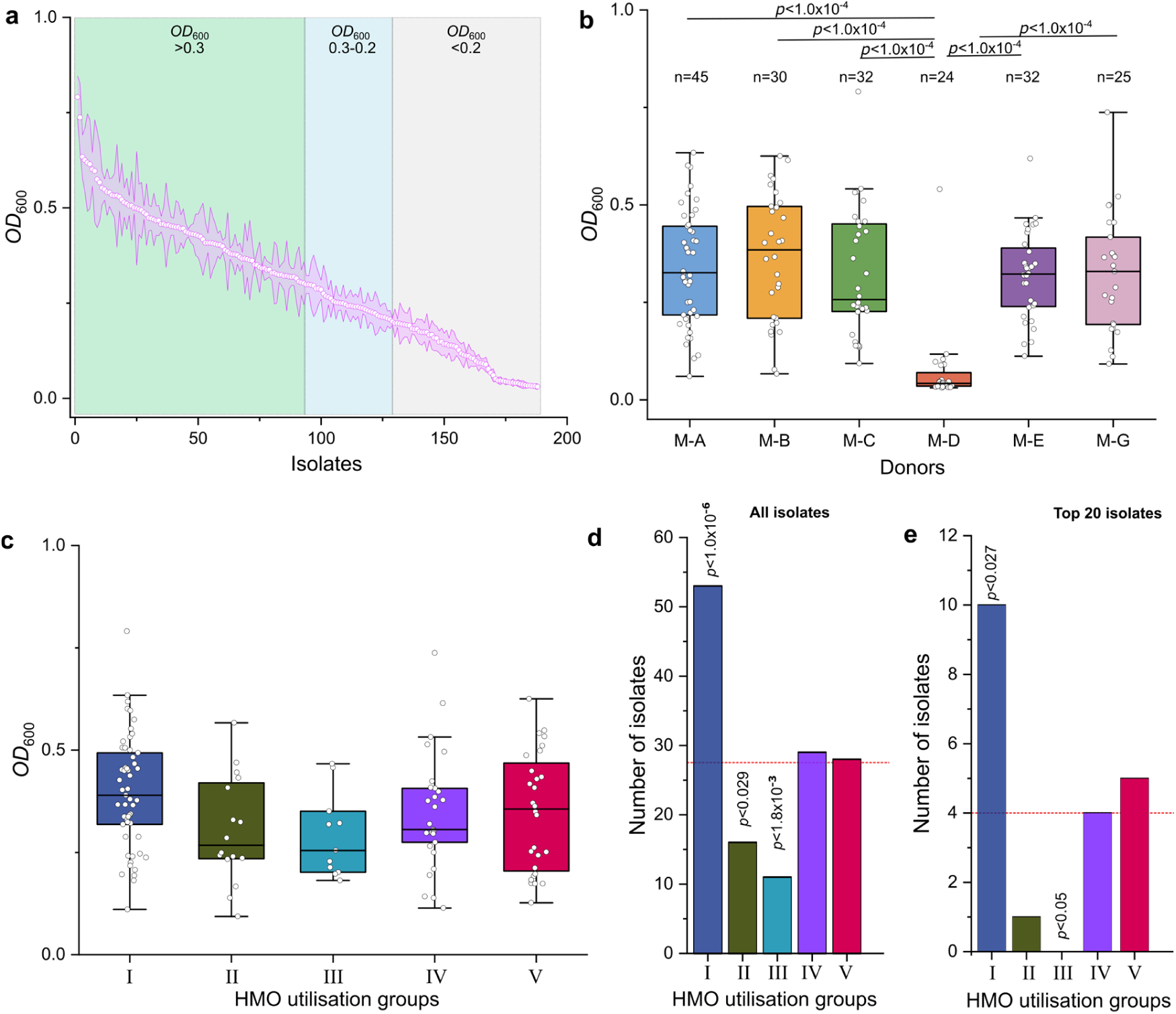
HMO utilisation profiles of maternal GM isolates. **a** Relative growth of adult mother GM isolates on HMOs compared to no-carbohydrate controls. The growth on HMOs, corrected for the growth on no-carbohydrate controls. Data represent averages (pink circle) from three independent growth experiments (n=3) with standard deviations (pink band). **b** Growth levels of isolates from different donors. Box plots show growth median values for isolates from each donor, with donor D exhibiting significantly lower growth levels compared to others. **c** Growth levels of different HMO utilisation clusters. Box plots illustrate the growth statistics of the isolates across the distinct HMO utilisation groups. **d** Comparison of the HMO-utilising clusters. Bar plots show the proportions of isolates assigned to different HMO utilisation clusters with group I, being significantly overrepresented and group III significantly underrepresented. **e** Comparison of the HMO-utilising clusters, similarly to **d**, but considering only the 20 isolates with highest growth levels across the clusters. **b** and **c** A one-way Anova was used to determine statistical significance between the individual groups and a Tukey’s range test was applied post hoc to identify significant different groups. **d** and **e** A Chi-Square (χ^2^) Goodness-of-Fit Test was used to assess significant deviations among the observed counts from an expected uniform using Origin Pro v10.0.0.154.

**Supplementary Fig. 13.**
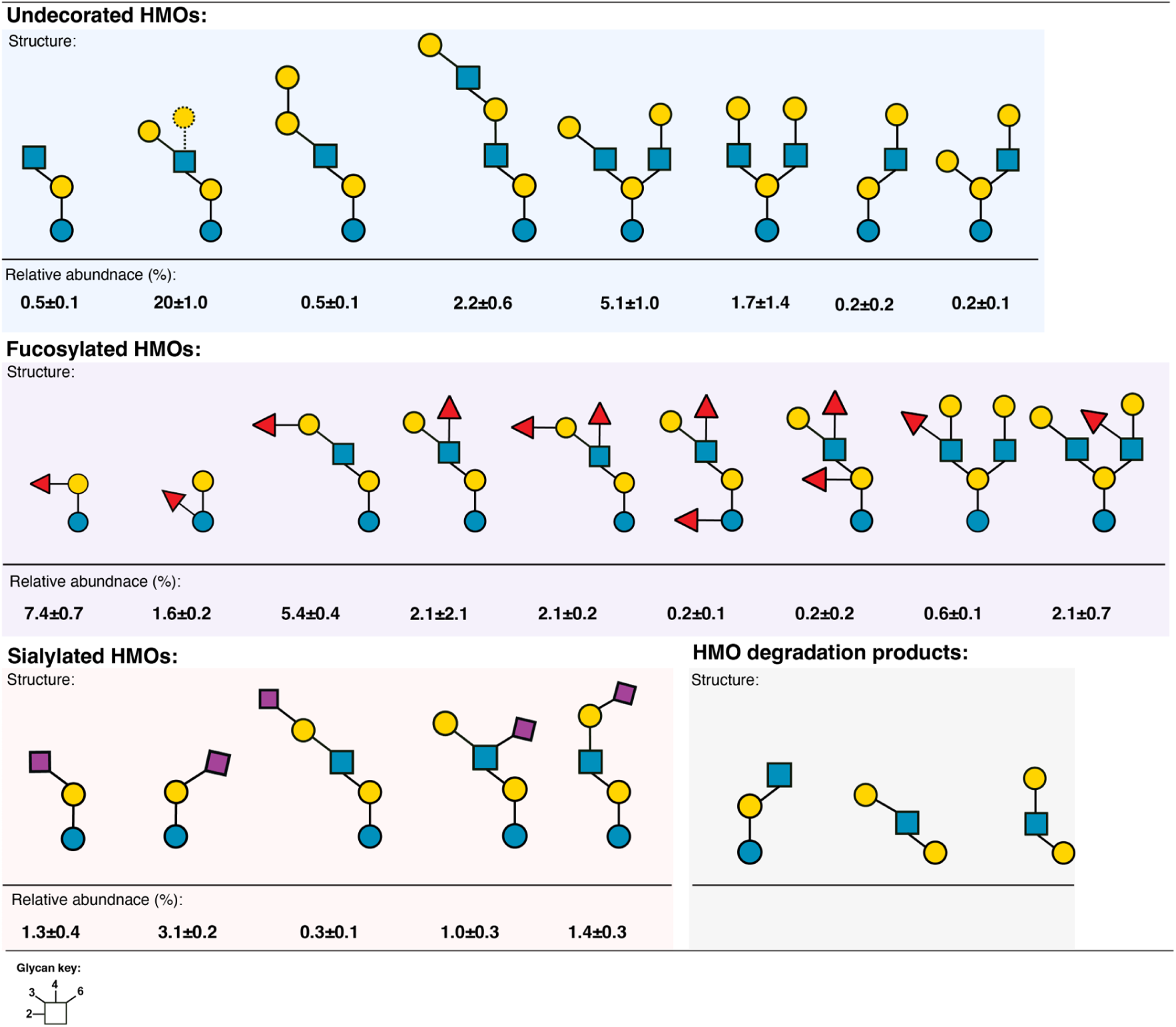
Relative abundance of HMOs selected for HMO depletion analysis. The reported glycan relative abundances are based on the area under the curve (AUC) of each structure normalised to the total AUC of 68 annotated structures in the HMO sample and expressed as a percentage. Data are represented as averages with standard deviations (SD) from three independent triplicates. The relative abundance of fucosyllactoses is lower than expected, due to losses during the removal of lactose from the HMO preparation. Glycan structures are presented according to the Symbol Nomenclature for Glycans (SNFG). The dashed line indicates the structural ambiguity between the typically more abundant LNT and less abundant LN*n*T due to their indistinguishability in the LC-MS/MS spectra^17^.

**Supplementary Fig. 14.**
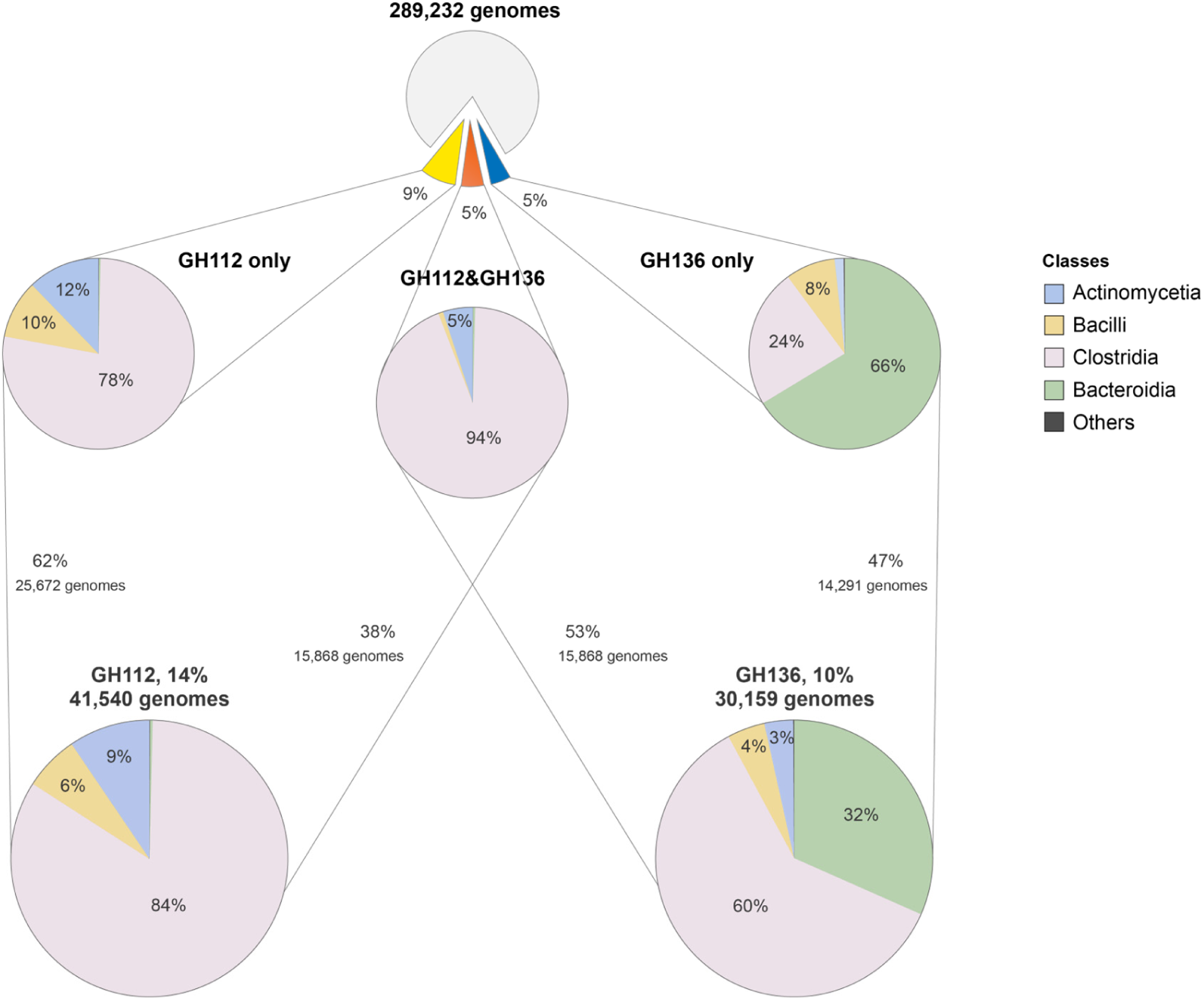
HMO-active GH112 and GH136 genes in the human GM catalogue^2^. Approximately 14% of the 289,232 genomes harbour GH112 homologues, with around 38% of these genomes encoding both GH112- and GH136 homologues. The GH136 homologues, including those co-occurring with GH112 and GH136, are harboured by 10% of the genomes in the catalogue. Classes with less than 1% abundance are not labelled in the figure.

**Supplementary Fig. 15.**
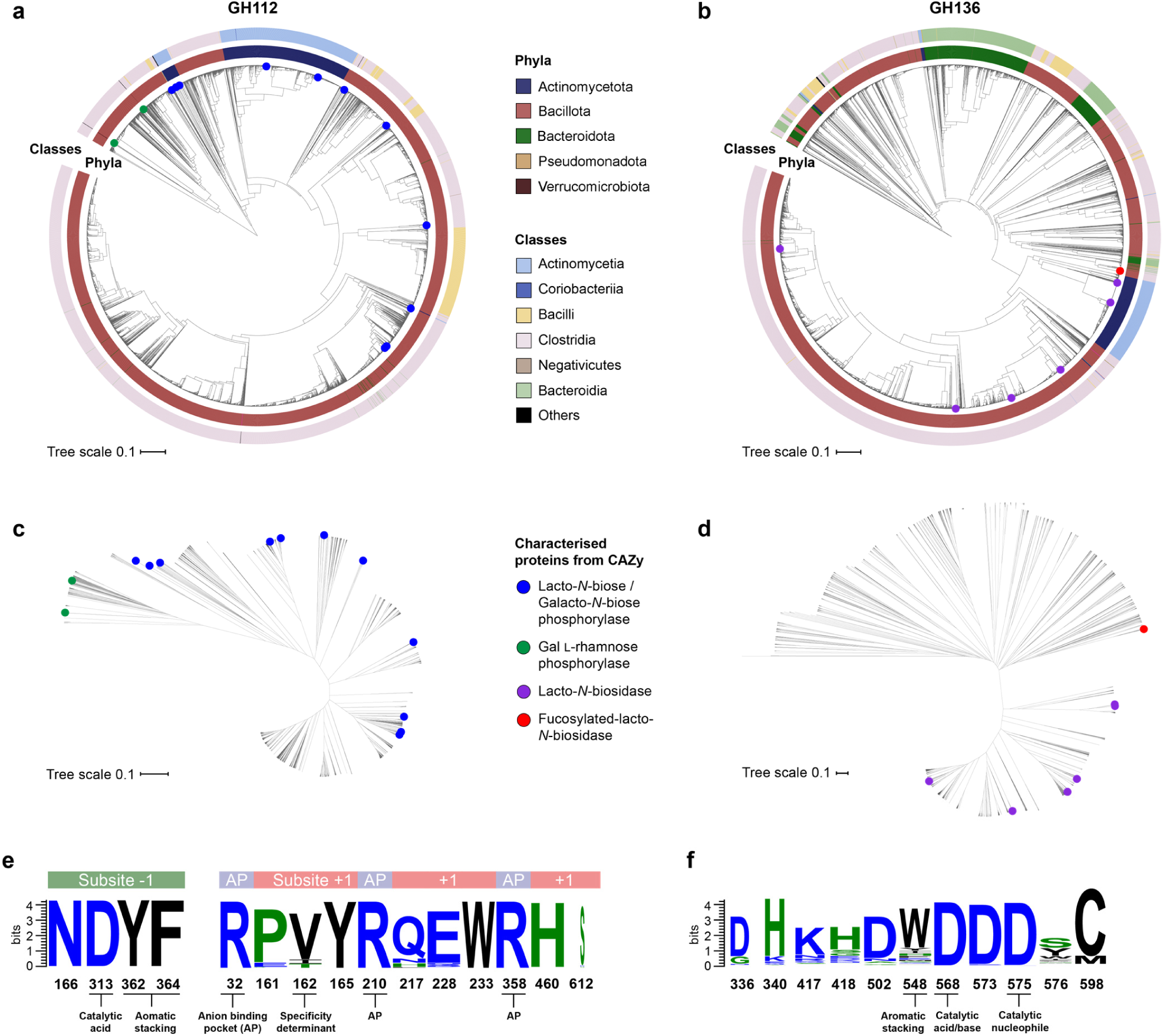
Phylogeny of GH112 and GH136, both mediating HMO degradation. **a** A total of 5,075 GH112 homologues and **b** 6,792 GH136 homologues retrieved from the human gut microbiome catalogue comprising 286,997 genomes. The taxonomic distribution of the sequences is colour-coded in the concentric circles. Characterised enzymes from the Carbohydrate-Active Enzyme (CAZy) database are marked as circles, colour-coded according to substrate specificities. **c** and **d,** Unrooted phylogenetic trees showing the evolutionary distances. **e** and **f** Sequence logos illustrating the conservation of both substrate binding subsites and catalytic residues in enzyme active sites, which supports the robustness of the annotation of retrieved sequences into these two GH families. The phylogenetic trees were generated using Interactive Tree of Life v6, and sequences logs were created with WebLogo 3.

**Supplementary Fig 16.**
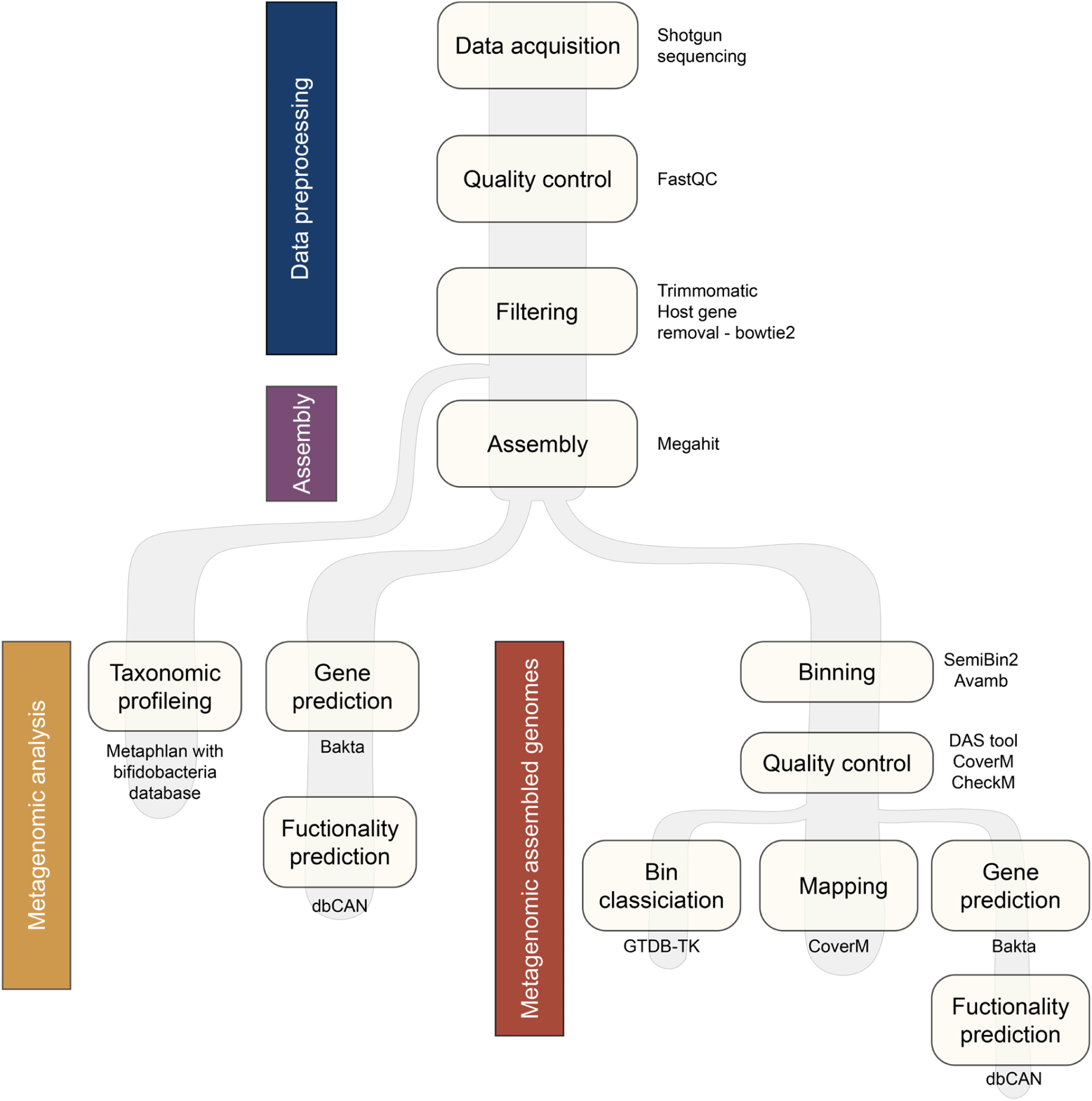
Metagenome sequencing, assembly and analyses pipeline. Whole shotgun metagenome (WSM) was performed on infant and mother’s faeces and enriched consortia.

### Supplementary Tables

**Supplementary Table 1.**
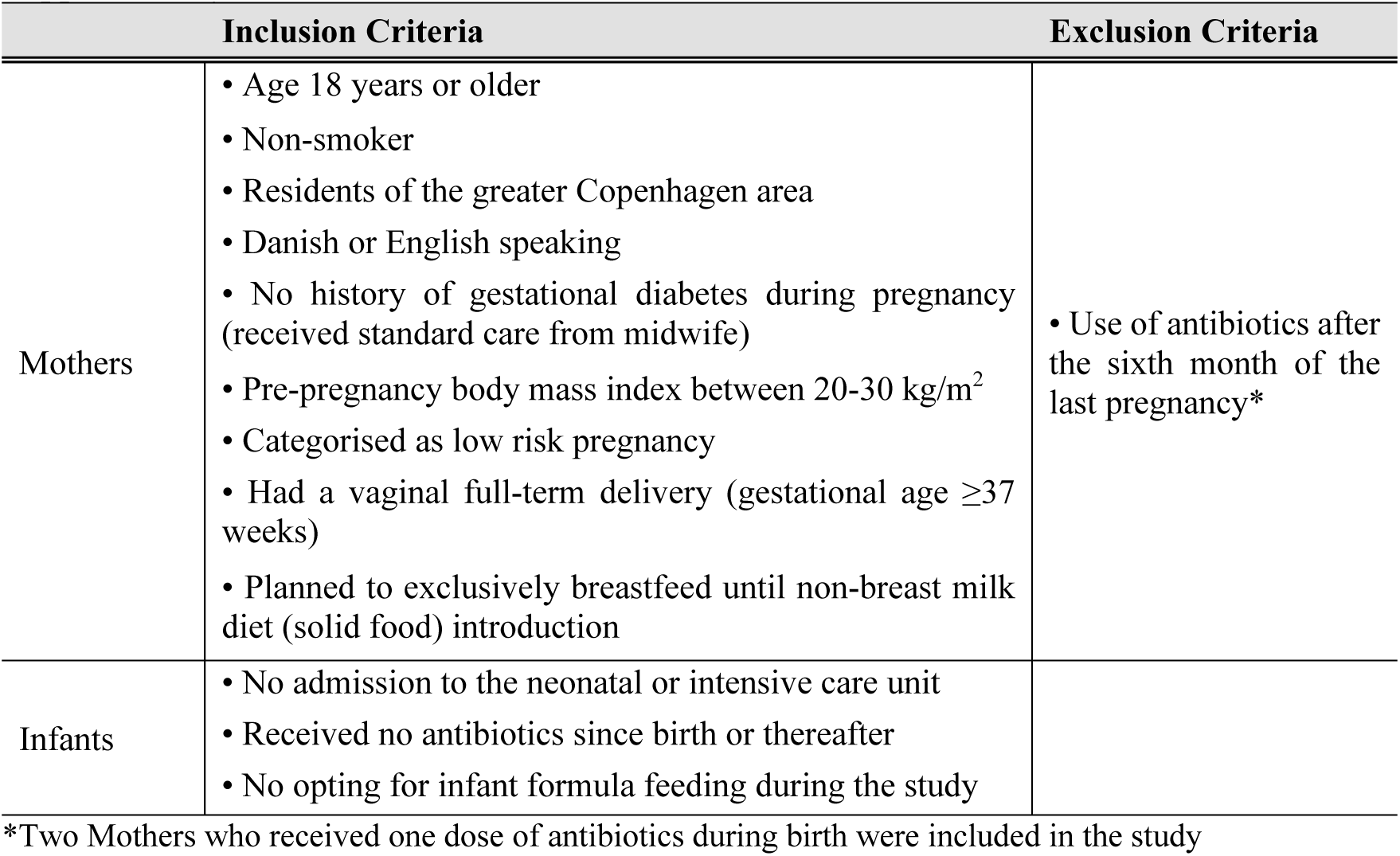
Inclusion and exclusion criteria for the Milkome cohort.

**Supplementary Table 2.** List of CAZy families specific to HMOs and host glycans. CAZymes are manually selected and curated data from www.CAZy.org^4^.

| Categories | Substrates | CAZYmes |
| --- | --- | --- |
| HMOs/host glycans | HMOs/host glycans | GH112 (Lacto-*N*-biose phosphorylase),<br>GH136 (Lacto-*N*-biosidase, Le<sup>a/b</sup> (fucosylated LNB) hydrolases),<br>GH20 (β-HexNAcase) |
|  | Fucosyl substitutions | GH29, GH95, GH151 |
|  | Sialyl substitutions | GH33, GH58, GH156, GH177, GH181 |
| HMOs/host* glycans | HMOs including LNT | GH42 (β-galactosidases) |
| Host glycans | Glycolipid | GH59 |
|  | Glycosaminoglycans (GAGs) | GH111, PL8_1, PL8_2, PL8_3, PL12, PL12_1, PL12_2, PL12_3, PL13, PL15_2, PL21, PL21_1, PL29, PL30, PL33, PL33_1, PL35 |
|  | *N*-glycans | GH5_18, GH163 |
|  | *O*-glycans | GH84, GH89, GH101, GH109, GH110, GH129 |

\*The GH42 CAZy family has clades which are specific for plant-derived galactosides, see SupplementaryData Fig. 6.

**Supplementary Table 3.**
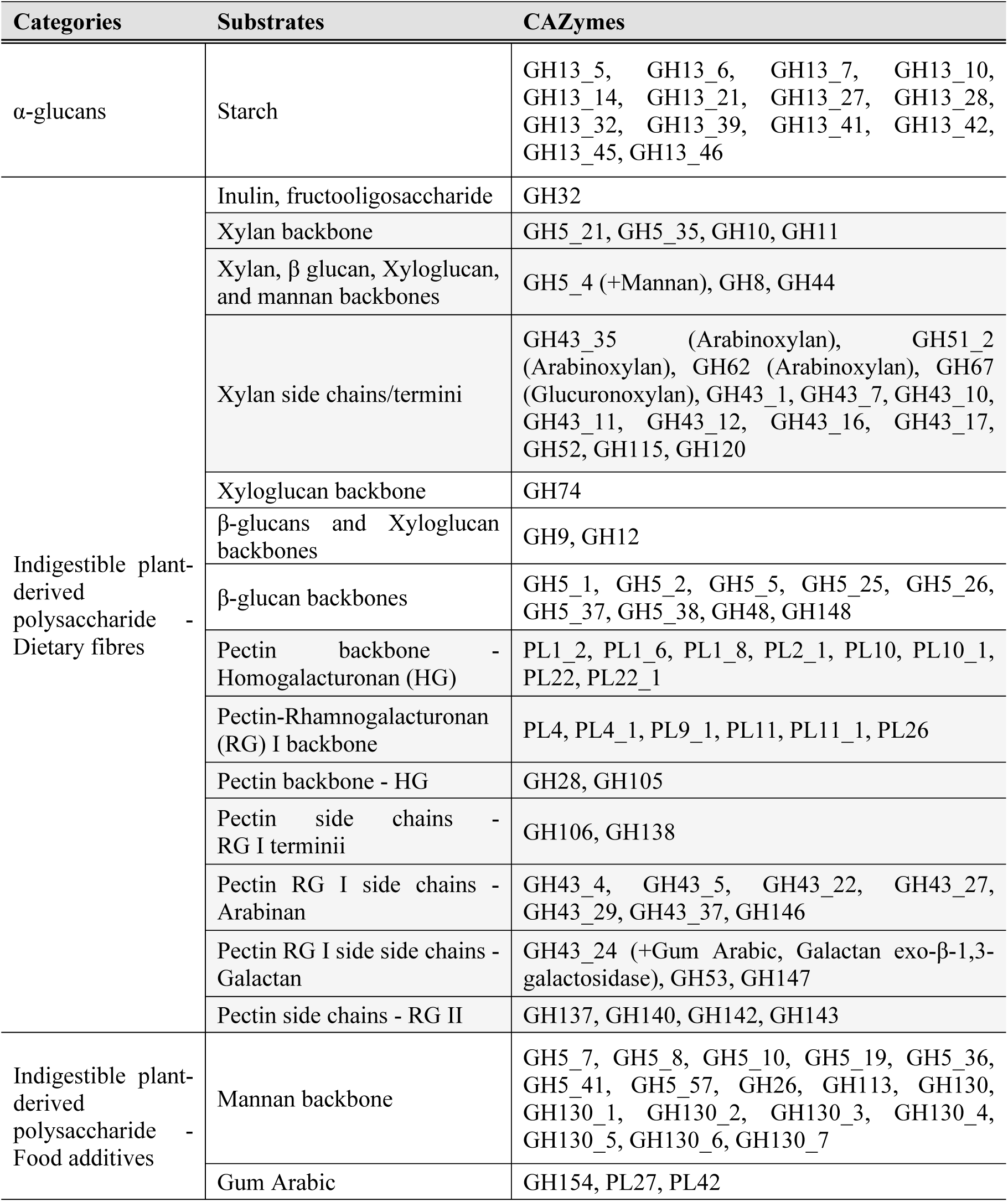
List of CAZy families specific to plant-derived polysaccharides. The CAZymes were manually selected and curated data from the CAZy database www.CAZy.org^4^

| Categories | Substrates | CAZymes |
| --- | --- | --- |
| α-glucans | Starch | GH13_5, GH13_6, GH13_7, GH13_10, GH13_14, GH13_21, GH13_27, GH13_28, GH13_32, GH13_39, GH13_41, GH13_42, GH13_45, GH13_46 |
| Indigestible plant-derived polysaccharide - Dietary fibres | Inulin, fructooligosaccharide | GH32 |
|  | Xylan backbone | GH5_21, GH5_35, GH10, GH11 |
| | Xylan, $\beta$ glucan, Xyloglucan, and mannan backbones | GH5_4 (+Mannan), GH8, GH44 |
|  | Xylan side chains/termini | GH43_35 (Arabinoxylan), GH51_2 (Arabinoxylan), GH62 (Arabinoxylan), GH67 (Glucuronoxylan), GH43_1, GH43_7, GH43_10, GH43_11, GH43_12, GH43_16, GH43_17, GH52, GH115, GH120 |
|  | Xyloglucan backbone | GH74 |
| | β-glucans and Xyloglucan backbones | GH9, GH12 |
| | β-glucan backbones | GH5_1, GH5_2, GH5_5, GH5_25, GH5_26, GH5_37, GH5_38, GH48, GH148 |
|  | Pectin backbone - Homogalacturonan (HG) | PL1_2, PL1_6, PL1_8, PL2_1, PL10, PL10_1, PL22, PL22_1 |
|  | Pectin-Rhamnogalacturonan (RG) I backbone | PL4, PL4_1, PL9_1, PL11, PL11_1, PL26 |
|  | Pectin backbone - HG | GH28, GH105 |
|  | Pectin side chains - RG I termini | GH106, GH138 |
|  | Pectin RG I side chains - Arabinan | GH43_4, GH43_5, GH43_22, GH43_27, GH43_29, GH43_37, GH146 |
| | Pectin RG I side side chains - Galactan | GH43_24 (+Gum Arabic, Galactan exo-β-1,3-galactosidase), GH53, GH147 |
|  | Pectin side chains - RG II | GH137, GH140, GH142, GH143 |
| Indigestible plant-derived polysaccharide - Food additives | Mannan backbone | GH5_7, GH5_8, GH5_10, GH5_19, GH5_36, GH5_41, GH5_57, GH26, GH113, GH130, GH130_1, GH130_2, GH130_3, GH130_4, GH130_5, GH130_6, GH130_7 |
|  | Gum Arabic | GH154, PL27, PL42 |

**Supplementary Table 4.** Media for bacterial enrichment and isolation.

| **Abbreviation** | **Designated use** | **Nitrogen source and mucin supplementation** |
| --- | --- | --- |
| YCFA | Reference medium | 0.25% (w/m) yeast extract<br>1% (w/m) Casitone |
| mYCFA | First enrichment of cohort samples | 0.1% (w/m) yeast extract<br>1% (w/m) Casitone<br>0.05% (w/m) Porcine mucin |
| mYmCFA | Second enrichment of cohort samples | 0.1% (w/m) yeast extract<br>0.5% (w/m) Casitone<br>0.05% (w/m) Porcine mucin |

**Supplementary Table 5.**
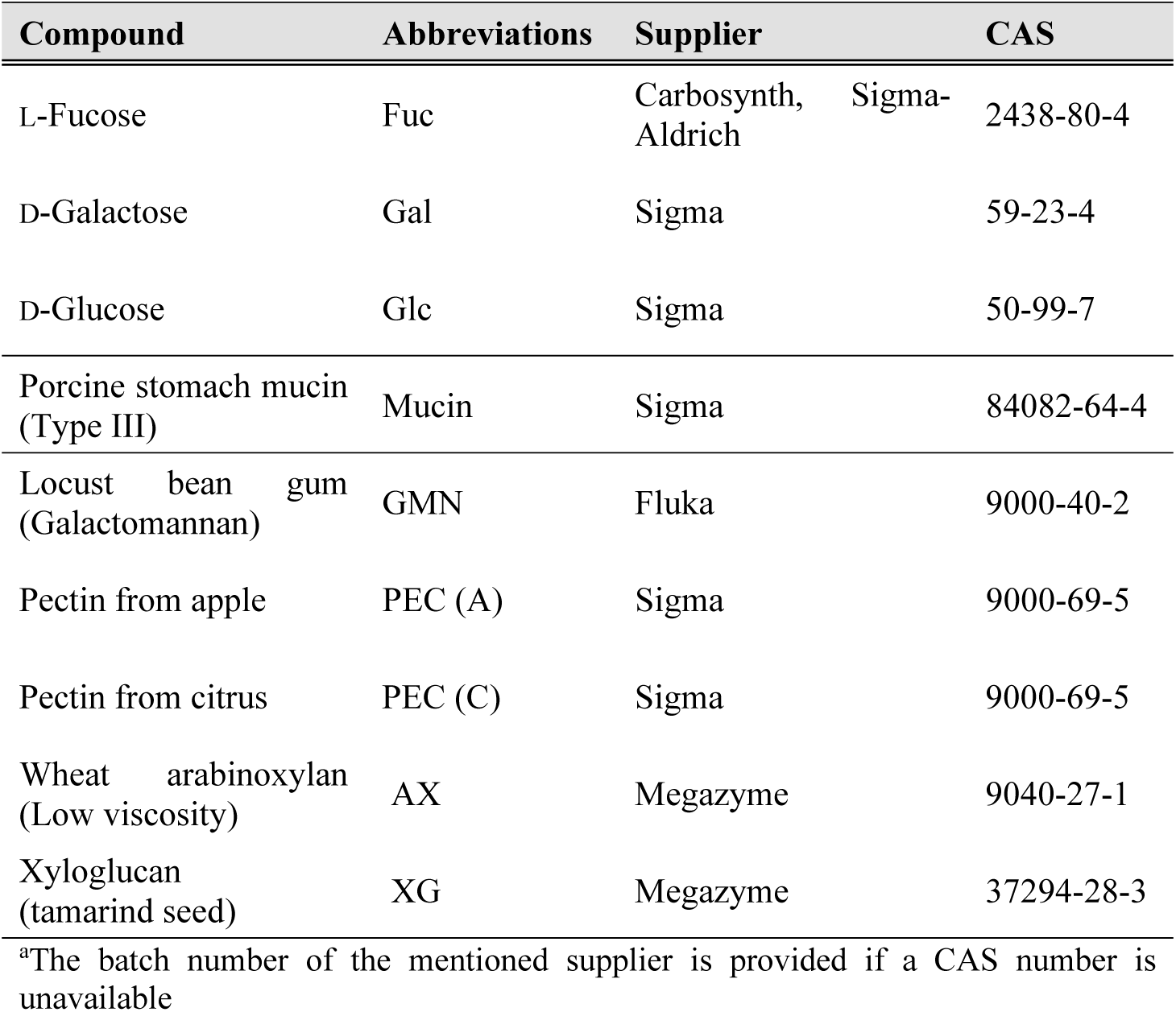
Carbohydrates substrates used in this study.

**Supplementary Table 6.** Substrates for enrichment of faecal samples.

| **Substrates for first enrichment (I)** |  |  |  |  |
| --- | --- | --- | --- | --- |
| **Compound** | **Content in medium (% w/v)** | **Infant samples** | **Mother samples** |  |
| HMOs | 0.5 | x | x |  |
| Porcine mucin | 0.5 | x | x |  |
| Arabinoxylan + Pectin (A) + Pectin (C) | 0.5 + 0.25 + 0.25 | x |  |  |
| Xyloglucan | 0.5 | x |  |  |
| Galactomannan + Gum Arabic | 0.5 + 0.5 | x |  |  |
| Glucose + Galactose + Fucose | 0.5 + 0.5 + 0.5 | x | x |  |
| **Substrates for second enrichment (II)** |  |  |  |  |
| **Starting consortia** | **Compound** | **Content in medium (% w/v)** | **Infant samples** | **Mother samples** |
| HMOs | HMOs | 1 | x | x |
| Porcine mucin | Porcine mucin | 0.5 | x | x |
| Arabinoxylan + Pectin (A, C) | Arabinoxylan | 0.5 | x |  |
| Arabinoxylan + Pectin (A, C) | Pectin (A, C) | 0.25 + 0.25 | x |  |
| Xyloglucan | Xyloglucan | 0.5 | x |  |
| Galactomannan + Gum Arabic | Galactomannan | 0.5 | x |  |
| Galactomannan + Gum Arabic | Gum Arabic | 0.5 | x |  |

**Supplementary Table 7.**
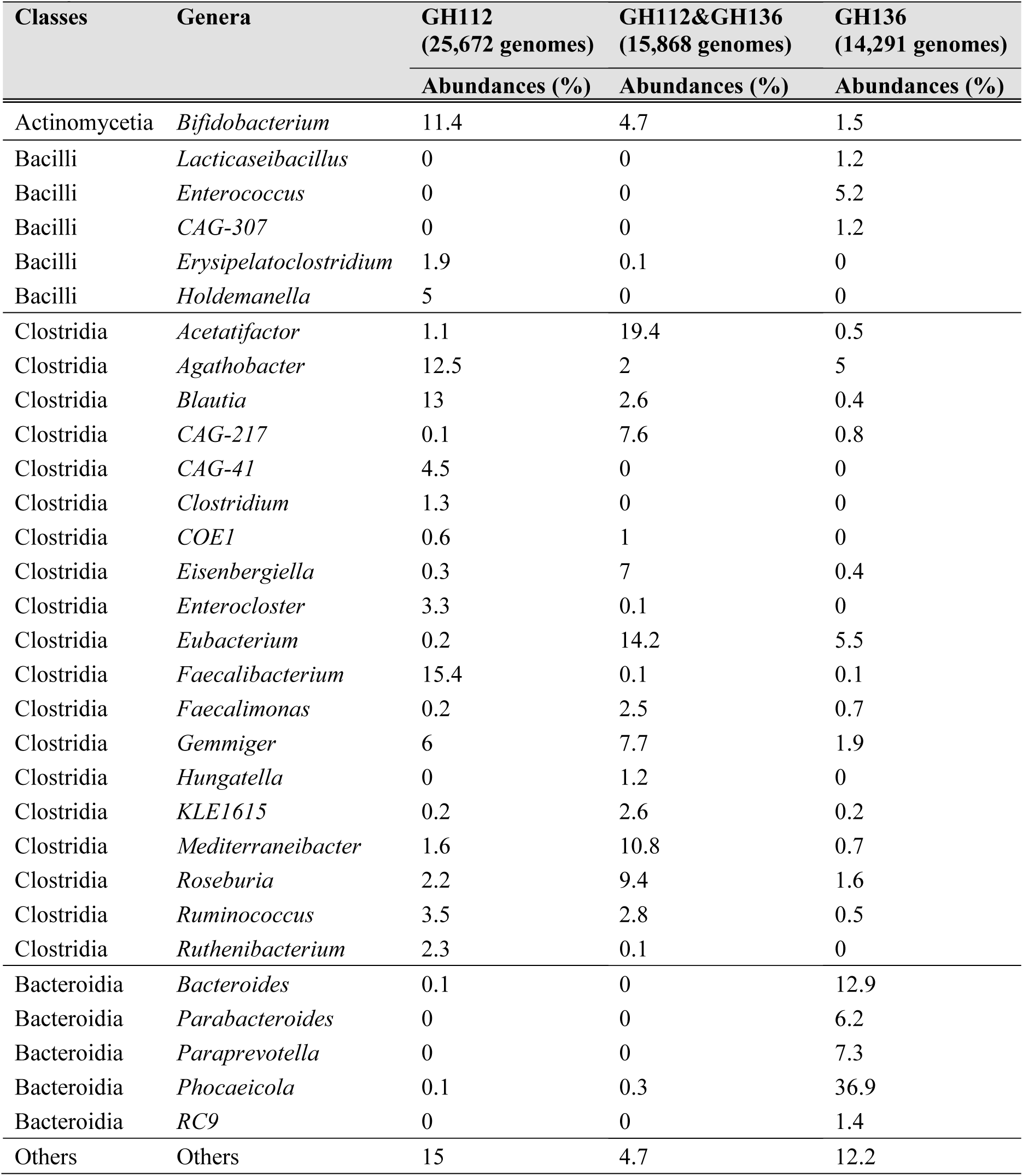
Abundances of GH112 and GH136 encoding genes in the HGM catalogue. ^2^.

